# Myosin-X sorts actin filaments into parallel bundles and tunes barbed-end dynamics

**DOI:** 10.64898/2026.09.08.749704

**Authors:** Wouter Kools, Sara Faour, Aliette Carlier, Trayana Hristova, Maria Elena Sirkia, Ludivine Chaix, Antony Lee, Pascal Martin, Anne Houdusse, Guillaume Romet-Lemonne, Antoine Jégou

## Abstract

Whether a molecular motor can spontaneously organize actin filaments into bundles has remained an open question despite decades of investigation. Here, we show that the dimeric myosin-X, essential for filopodia initiation and extension in cells, is capable of sorting actin filaments into parallel bundles, gathering barbed ends within 1-2 µm. We observe that myosin-X processivity is comparable on single filaments and on bundles induced by myosin-X or fascin. Upon reaching barbed ends, myosin-X slows down the addition or removal of actin subunits, in a myosin density-dependent manner. Furthermore, the funneling of myosin-X towards the remaining filaments at the bundle tip increases motor density and triggers dynamic clustering. Together, we propose that the motor activity of myosin-X is sufficient to initiate filopodia, independently of passive crosslinkers such as fascin or fimbrin.

**Summary:** Myosin-X, a processive dimeric motor, crosslinks and sorts actin filaments based on their polarity to create parallel actin bundles. It accumulates at bundle tips and tunes barbed-end actin elongation.

## Introduction

Filopodia are dynamic cell projections that drive the exploration of, and communication with, their environment (Blake and Gallop, 2023; Jacquemet et al., 2015). They are flexible structures, displaying alternating phases of elongation and retraction, with lifetimes on the order of a few minutes. Filopodia have been observed in many cell types and at virtually all stages of organismal development (e.g. dorsal tube closure in Drosophila, neurogenesis,…). Through cell-surface receptors such as integrins and cadherins, filopodia integrate signals from various extracellular cues (Blake and Gallop, 2023; Jacquemet et al., 2015). They are built by the simultaneous growth of 10–30 actin filaments organized into parallel bundles, with their barbed ends oriented towards the filopodial tip (Faix and Rottner, 2006), and extend at a velocity of a few hundred nm/s (Bornschlögl, 2013).

Myosin-X, an unconventional dimeric myosin motor, is by far the most abundant molecular motor in filopodia, where it strongly accumulates at the tip (Berg et al., 2000; Tokuo and Ikebe, 2004). It belongs to the MyTH4-FERM myosin subfamily, together with myosins-VII and -XV, which all derive from a single ancestral myosin present in the last common ancestor of holozoans (Sebé-Pedrós et al., 2014). This subfamily has been proposed to have emerged alongside the increasing complexity of actin networks in unicellular holozoans, prior to the advent of multicellularity (Houdusse and Titus, 2021; Sebé-Pedrós et al., 2014). Myosin-X mediates the attachment of actin bundles to the plasma membrane in filopodia, either directly, through its PH domain binding PIP3 lipids, or indirectly, through MyTH4 and FERM domain binding partners that are themselves membrane-associated (Houdusse and Titus, 2021).

Myosin-X has long been recognized as one of the factors required to initiate and maintain filopodia extension (Berg et al., 2000; Bohil et al., 2006; Fitz et al., 2023; Pokrant et al., 2023; Tokuo et al., 2007), and it was quickly established that its motor function, rather than its cargo-binding function, is critical for filopodia formation (Tokuo et al., 2007). Dimeric myosin-X walks processively on actin filaments (Bao et al., 2013; Ropars et al., 2016; Sato et al., 2017; Sun et al., 2010), with a high duty ratio similar to that of the well-studied myosin-V, that is, a single myosin-X motor domain spends a large fraction of its ATPase cycle bound to actin (Homma and Ikebe, 2005). In addition, myosin-X harbors an anti-parallel coiled-coil for dimerization that appears to be unique among myosins (Lu et al., 2012; Ropars et al., 2016), together with an extended lever arm thought to be more flexible than that of myosin-V, since it contains only 3 IQ motifs (calmodulin-binding sites) that do not cover its full length (Knight et al., 2005). Owing to this combination of an extended lever arm and an anti-parallel coiled-coil, myosin-X displays a multi-modal distribution of step sizes, with steps as large as 50 nm (Nguyen et al., 2023; Ropars et al., 2016). The contribution of this coiled-coil to myosin-X’s walking properties (velocity, run length, step sizes, …) and to its specificity for fascin-actin bundles has been examined in several studies (Bao et al., 2013; Caporizzo et al., 2018; Nagy et al., 2008; Ropars et al., 2016; Sato et al., 2017; Sun et al., 2010), sometimes with observations that are hard to reconcile. Notably, analysis of myosin-X step sizes on fascin-actin bundles indicates that myosin-X frequently side-steps onto neighboring filaments, whose spacing and registration are strongly imposed by fascin (Aramaki et al., 2016; Gong et al., 2025). Because the stepping of myosin dimers requires the trailing motor domain to detach only once the leading motor domain has bound actin (Baboolal et al., 2016; Sakamoto et al., 2008), side-stepping on fascin-actin bundles means that a single myosin-X dimer actually crossbridges two actin filaments during part of the ATPase cycle of each motor domain.

At early stages of filopodia formation, actin filaments need to be organized into parallel arrays, and the actin elongator Ena/VASP is recruited to the initiation foci for filopodia elongation to proceed (Pokrant et al., 2023). Filopodia initiation appears to occur prior to the recruitment of filopodia-specific crosslinking proteins such as fascin (Svitkina et al., 2003). Yet, despite myosin-X’s specificity for filopodia in vivo and for fascin-actin bundles in vitro, how myosin-X may organize actin filaments during filopodia initiation remains unknown. Similarly, whether this process requires additional proteins, such as Ena/VASP or RAPH1/Lamellipodin is also not fully resolved.

Here, using in vitro reconstitution, we show that myosin-X alone is sufficient to sort actin filaments by their polarity and organize them into parallel bundles with close-by barbed ends. We dissect the molecular events underlying this filament organization. We further report that this organization permits to concentrate myosin-X motors at the tip of bundles, favoring the formation of dynamic myosin clusters. In turn, the increase in motor density slows down barbed-end assembly dynamics. We propose that, in cells, myosin-X motor activity is sufficient to initiate parallel actin bundles, prior to the recruitment of the fascin crosslinker that subsequently drives filopodia elongation.

## Results

### Myosin-X creates parallel actin bundles and accumulates at their tip

To investigate the ability of myosin-X to initiate filopodia extension, we studied the bundling of actin filaments by myosin-X using a series of *in vitro* assays. Following the rationale from Ropars et al (Ropars et al., 2016), experiments were performed using a human myosin-X construct containing the motor domain, the lever arm, the anti-parallel coiled-coil, followed by a 19 amino-acid linker and a GCN4 leucine zipper to keep the motor dimerized, but lacking the C-terminal cargo-binding domains (see Methods). This construct carries an mStayGold at the C-terminus, reasoning that a fluorescent protein at its N-terminus could sterically interfere with the binding interface between the motor domain and the actin subunits of the filament. We confirmed that purified myosin-X were mostly dimers, based on the fluorescence intensity of isolated myosins walking processively on filaments (supp. Fig.1). All experiments were performed at 25°C, pH 7.4, 50 mM KCl, 1 mM MgCl_2_, and different concentrations of Mg-ATP and Mg-ADP (see Methods).

In a surface-passivated in vitro chamber assay, pre-assembled actin filaments (average length ∼ 5 µm, not stabilized by phalloidin) were first injected and subsequently exposed to a few nanomolar of myosin-X, in a buffer containing both high concentrations of ATP and ADP, supplemented with methylcellulose (Fig. 1A, B). On its own, without myosins, the presence of methylcellulose at our concentrations does not induce filament bundling (Fig. 1A). With myosin-X, actin bundles formed within a few seconds, with parallel filaments, as reported by the walking direction of myosins (Fig. 1C), and by the fact that barbed ends were mostly localized at one bundle tip only (Fig. 1E). Bundle size was on average 16 ± 12 filaments (n = 41 bundles, at 200 µM ATP, 400 µM ADP), with barbed ends gathered within 1-2 µm (Fig. 1D, Supp. Fig. S3).

**Fig. 1.**
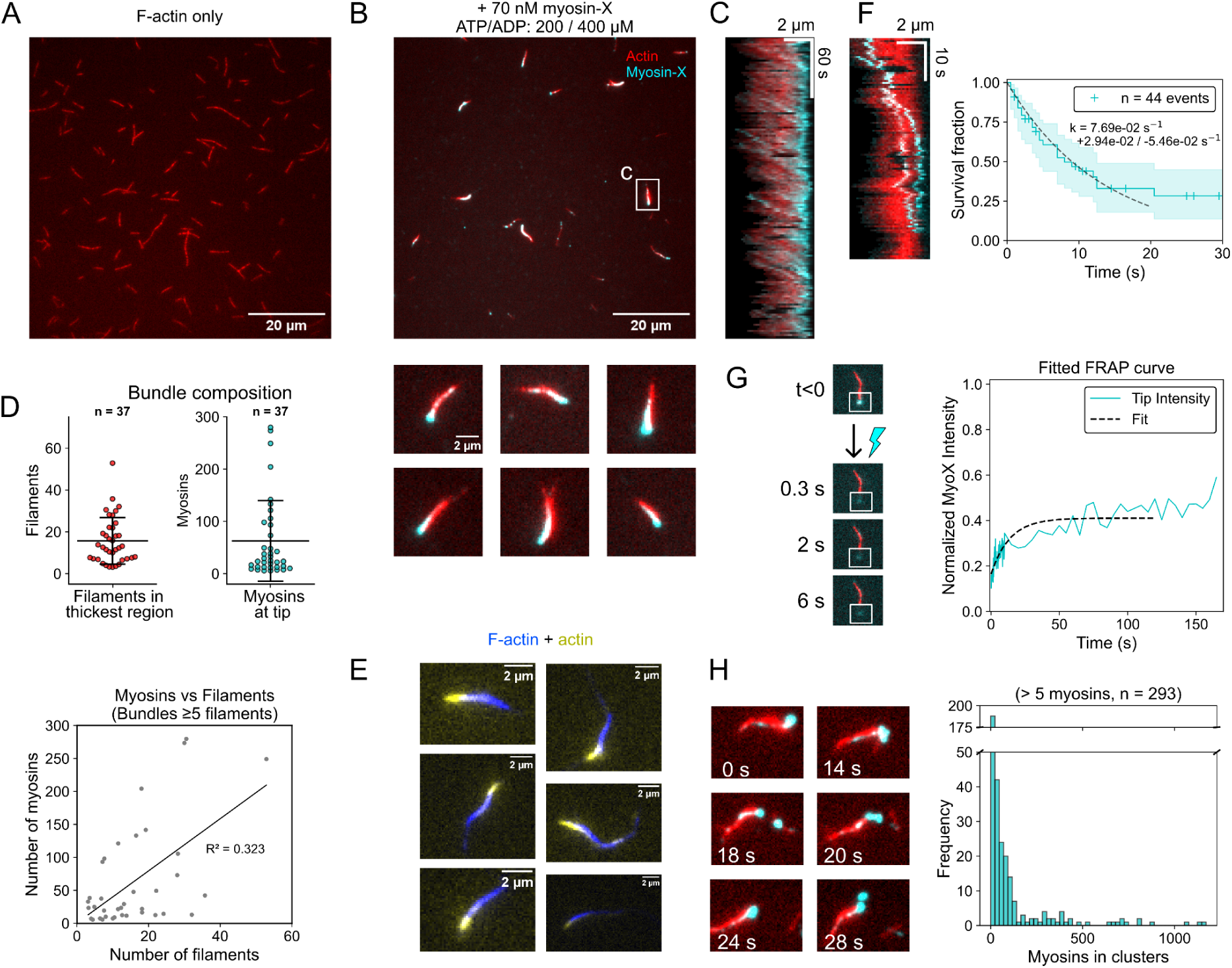
Myosin-X organises actin filaments into parallel bundles and accumulates at bundle tips. (A) Pre-polymerized 15% Alexa568-labeled actin filaments (0.1 µM) in an open-chamber assay, in F-buffer, in the presence of 200µM ATP and 0.2% methylcellulose. Filaments do not form bundles. (B) Top: After the addition of 70 nM dimerised myosin-X, in the presence of 200 µM ATP, 400 µM ADP and 0.2 µM unlabelled G-actin, bundles spontaneously form within seconds. Scale bar: 20 µm. Bottom: Six examples of bundles from the same experiment. Scale bar applies to all examples. We verified that myosin-X activity was not strongly affected by the presence of methylcellulose (see Methods, Supp. Fig. S15). (C) Both ends of the boxed bundle in B were tracked manually using TrackMate. The tip where myosin accumulates was chosen as a reference and aligned to create the kymograph. (D) Top: bundle thickness and number of myosins accumulated at bundle tip. Error bars are standard deviations. Data from two independent experiments at 200 µM ATP and 400 µm ADP. Bottom: number of myosins as a function of number of filaments. (E) In an experiment as described in B, with unlabeled myosin-X, 0.25 µM 15% Atto643-labeled G-actin was added to reveal barbed end location in bundles composed of pre-polymerized 8% Atto488-labelled actin filaments. Scalebar: 2 µm. (F) Left: Kymograph of a bundle from an experiment with only 0.2% myosin-X labelled with mStayGold, shows myosins walking toward the bundle tip and dwelling there. Right: Survival fraction of myosin-X dwelling at bundle tips. Fit of the survival curve by a single exponential decay function (k_off_ = 0.077 s^-1^, n = 44, from 1 experiment). (G) Recovery after photobleaching of the fluorescence intensity of accumulated myosin-X at a bundle tip. (H) Left: Example of myosin-X clusters separating and fusing over time. Right: Distribution of the number of myosin-X in clusters (n = 200, from two independent experiments).

Having both ATP and ADP present in solution was chosen as myosins form tighter bundles that are thus easier to observe and follow over time. Moreover, physiological ADP concentration is elevated and almost equimolar to ATP at the cell periphery (Holland and Gallo, 2023; Schuler et al., 2017; Tantama et al., 2013). Still, bundles could form at saturating ATP concentrations in the absence of ADP with a sufficient concentration of myosins (Supp. Fig S2). Besides, experiments with longer actin filaments led to contractile actin meshworks, in which the high connectivity induced by myosin-X prevented reorganization into isolated parallel bundles (Supp. Fig. S4).

Interestingly, the myosin-X distribution revealed that motors accumulated at the tip of the bundles (average = 63 myosins, Fig. 1D), consistent with a ‘funneling’ effect, where the most advanced barbed ends of a bundle would collect motors from the trailing filaments. On some occasions, myosin-X at bundle tips formed large puncta (Fig. 1H). These large puncta can be classified as clusters, as they eventually detached from bundle tips while remaining cohesive as they diffused away in solution (Supp. movie 1). Myosin-X clusters formed only in the presence of actin filaments, as myosin-X alone in solution did not cluster. Tracking of individual myosins revealed that they dwell at bundle tips, for ∼ 9 seconds, likely integrating a cluster (Fig. 1F). Fluorescence recovery after photobleaching (FRAP) of myosin clusters at bundle tips showed that half of the myosin population turned over, providing support for the “cohesiveness” of the myosins at bundle tips (Fig. 1G). The recovery of the mobile fraction (t_1/2_ ∼ 4 seconds, Fig. 1G) is faster than the steady-state flux of myosins along the bundle could account for (see below). This indicates that at least part of the myosin population in the clusters dynamically exchanges with myosins in solution. Consistently, the early phase of the fluorescence recovery of the myosin intensity at bundle tip was similar when myosins along the bundle shaft were also photobleached (Supp. Fig. S5).

Overall, these observations show that myosin-X spontaneously organizes filaments into parallel actin bundles with gathered barbed ends, leading to a sharp increase in motor density at bundle tips.

### Myosin-X accommodates processive stepping on various actin organizations

We next investigated which biochemical characteristics of myosin-X are responsible for bundle formation. First, we quantified the affinity of our myosin-X construct for ATP. We measured the velocity of single myosin-X molecules on individual actin filaments, anchored by their pointed end on the surface of a microfluidics chamber and aligned with the flow (Fig. 2A). In this assay, filaments are on average ∼ 200 nm above the surface to avoid any artifact (Wioland et al., 2020), and filaments were not stabilized by phalloidin. We recently showed that a small viscous drag force, applied by the flow on individual walking myosin-X, does not impact their velocity (Bagès et al., 2025). Quantifying the myosin-X velocity as a function of ATP concentration (Supp. Fig. S6) allowed us to estimate both the ATP-dependent (k_on, ATP_ = 13.5 (± 5) µM^-1^.s^-1^) and ATP-independent (k_off, ADP_ = 34 (± 4.5) s^-1^) parts of the ATPase cycle, in agreement with previous observations (Bao et al., 2013; Ropars et al., 2016; Sun et al., 2010). We obtained a maximum velocity of 1500 nm/s on single actin filaments at 25°C. This value is substantially larger than what has been reported previously by other labs (Caporizzo et al., 2018; Nguyen et al., 2023; Ropars et al., 2016; Sun et al., 2010; Takagi et al., 2014), and could originate from various factors (dimerisation strategy, absence of fluorescent reporter attached to the motor domain, …). Increasing ADP concentration in the presence of 200 µM ATP reduces myosin-X velocity on single actin filaments, with an apparent inhibition affinity of K_+ADP_ = 44 (± 17) µM (Supp. Fig. S6).

**Fig. 2.**
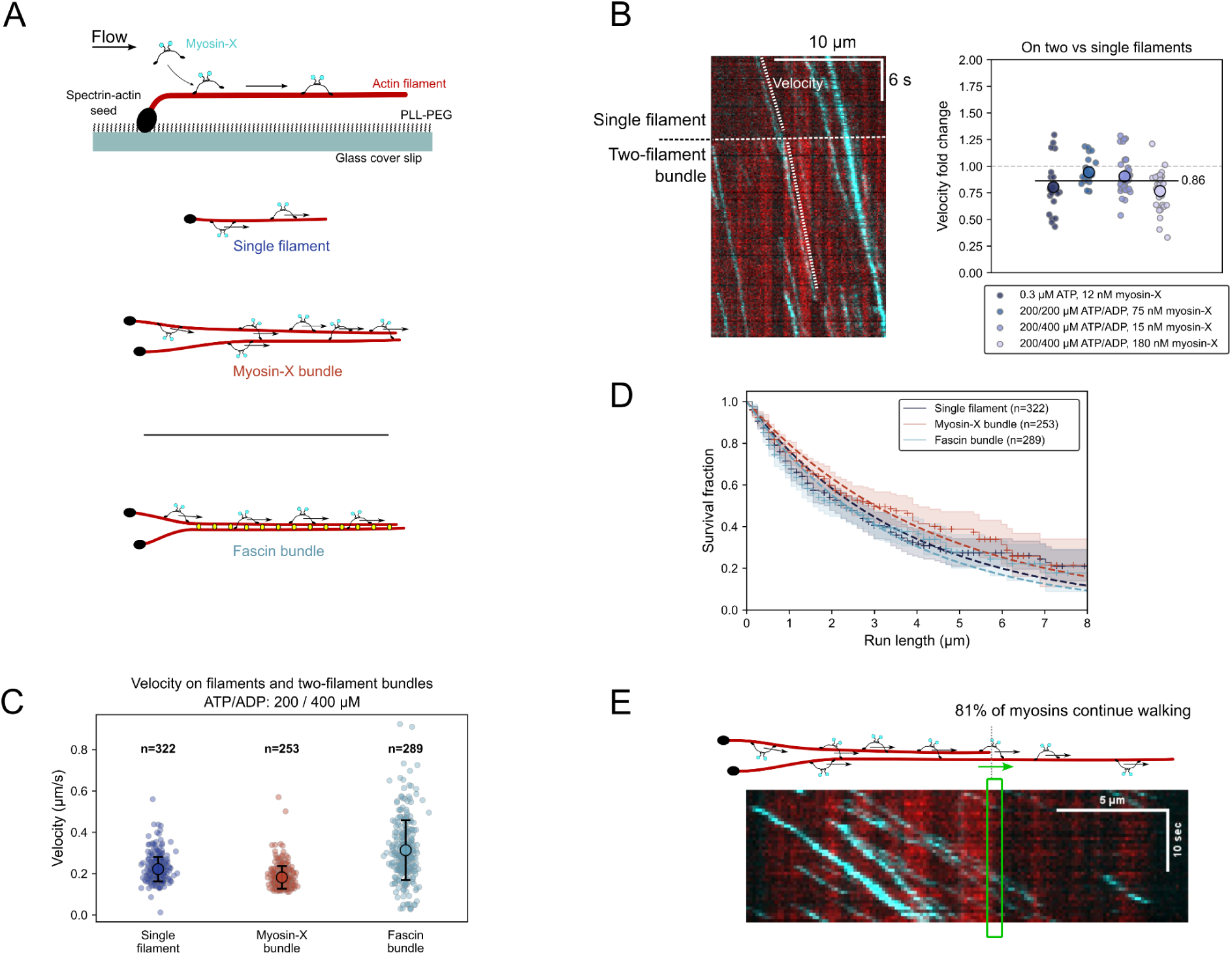
Walking behaviour of myosin-X on actin filaments and bundles. (A) Microfluidics experimental setup. In a microfluidic chamber, actin filaments are elongated from surface-anchored spectrin-actin seeds. The filaments align with the flow direction, on average 200 nm above the passivated surface. Two filaments in close proximity can form a bundle in the presence of myosin-X motors. Alternatively, filaments can first be cross-linked by 1 µM fascin, and then exposed to myosin-X and the same concentration of fascin. (B) The velocity of individual myosin-X on 2-filament myosin-X bundles compared to the velocity of the same myosins on single filaments (0.3 µM ATP, 12 nM myosin-X (n = 19); 200/200 µM ATP/ADP, 75 nM myosin-X (n = 26); 200/400 µM ATP/ADP, 15 nM myosin-X (n = 42); 200/400 µM ATP/ADP, 180 nM myosin-X (n = 7)). Large data points represent the average of each condition, and the horizontal line is the global average (see also Supp. Fig. S9). (C) Velocity of individual myosin-X on single filaments (n=322), 2-filament bundles induced by myosin-X (n=253) or by fascin (n=289), in the presence of 200 µM ATP and 400 µM ADP. Different conditions were acquired on the same day. Large data points represent the average of each condition. Error bars are standard deviations. At 200 µM ATP, myosin-X also walked more slowly on myosin-X bundles compared to single filaments (Supp. Fig. S7). (D) Survival fraction of myosin-X run length on individual myosin-X on single filaments (n=322), 2-filament bundles induced by myosin-X (n=253) or by fascin (n=289), as a function of time. Survival curves fitted with a single exponential decay function, gives a characteristic run length of 3.80 µm (95% CI: 2.99 - 4.84 µm) on single filaments, 4.5 µm (95% CI: 3.45 - 6.02 µm) on myosin-X-induced bundles, and 3.53 µm (95% CI: 2.77 - 4.55 µm) on fascin-induced bundles. Corresponding processivity data shown in Supp. Fig. S10. (E) Myosin-X detachment at the transition between 2 to 1 filaments of 2-filament bundles induced by myosin-X. 81% (± 7.5)% of myosins detach. (0.3 µM ATP and 12 nM myosin-X (n = 48 switching events, 77.3% detach), 200/200 µM ATP/ADP and 75 nM myosin-X (n = 82, 86.3% detach), 200/400 µM ATP/ADP and 15 nM myosin-X (n = 57, 72.2% detach), 200/400 µM ATP/ADP and 180 nM myosin-X (n = 29, 88.1% detach)).

In the presence of myosin-X, filaments in close proximity readily formed bundles in the microfluidics assay (Supp. Fig. S7), similar to what we observed in an open chamber (Fig. 1). We note that here there is no methylcellulose, showing that the presence of a crowding agent is not required. Comparing the velocity of myosin-X on single filaments and on 2-filament bundles induced by myosin-X, observed side by side in a microfluidics chamber, we found that the velocity of myosin-X was on average 16% lower on bundles than on single filaments (Fig 2B, Supp. Fig. S7). In comparison, the velocity of our myosin-X construct on fascin-induced 2-filament bundles was ∼ 10% faster than on single filaments (Fig. 2C, Supp. Fig. S8). As a comparison, full length mWasabi-myosin-X was reported to be twice as fast on large fascin-actin bundles than on single filaments (Ropars et al., 2016).

In the presence of 200 µM ATP and 400 µM ADP, the run length of myosin-X was 3.8 (± 0.34) µm on single actin filaments (n=322), 3.5 (± 0.30) µm on fascin-induced 2-filament bundles (n=289), and 4.5 (± 0.44) µm on myosin-X-induced 2-filament bundles (n=253) (Fig. 2D). Combined with measured velocities, these observations indicate that myosin-X is more processive on the loose bundles it induces itself. Compared to rigid structure of fascin-actin bundles (Claessens et al., 2006; Gong et al., 2025), the spacing between actin filaments in myosin-X-actin bundles likely fluctuates strongly, owing to the flexibility of the myosin-X lever arm (Lu et al., 2012; Ropars et al., 2016), and to the distance between two consecutive myosins-X crosslinking filaments (∼ 1 µm in our assay). Assuming that finding the next actin binding site is not rate-limiting during the myosin ATPase cycle, myosin-X-actin bundles seem thus to offer more accessible binding sites for forward stepping, but the larger inter-filament distance is expected to result in smaller displacements along the bundle’s long axis (Supp. Fig. S10).

Taken together, these results show that myosin-X is similarly processive on different actin architecture, and readily accommodates processive stepping on neighboring parallel filaments, only mildly impacting longitudinal velocity and run length.

### Myosin-X ‘funneling’ and accumulation at bundle tips

We next sought to investigate what causes myosin-X accumulation at bundle tips. End-pausing of processive motors at filament end is required for motor jamming on individual filaments, as reported for microtubules (Bieling et al., 2010; Leduc et al., 2012; Varga et al., 2009). Contrary to what we observed for individual myosin reaching bundle tips (Fig 1F), we did not detect any end residency of myosin-X at the barbed end of single actin filaments (see below).

‘Funneling’ could be another mechanism increasing motor density at bundle tips, whereby motors reaching the end of a filament continue walking on the other filaments down the bundle. The simplest, but also the most extreme, ‘funneling’ case is the transition from 2 to 1 filament. In this situation, 81 (± 7.5)% (N=4, n=194 motors) of the individual motors continue walking through the two-to-one filament transition segment (∼ 500 nm, as per our microscope resolution) (Fig. 2E). This percentage is lower, but compatible, with the 89% of motors that keep on walking over a 500 nm segment measured along 2-filament bundles (run length = 4.5 µm, Fig. 2D). Myosin-X thus efficiently steps onto a downstream filament of a myosin-X-actin bundle, which would increase motor density as the number of remaining filaments decrease towards the tip of a bundle.

### Myosin-X sorts filaments based on their polarity and is a stable crosslinker

To assess the ability of myosin-X to create parallel actin bundles, we examined the pairing of 2 filaments by myosin-X in an open-chamber assay with only very few filaments freely diffusing (limited to a 2D diffusion by the presence of the crowding agent). To identify filament polarity, barbed ends were elongated from actin fluorescently labeled in a different color than the filament core (Supp. Fig. S11, Supp. movie 2). As expected, anti-parallel filaments quickly slid apart, while parallel filaments remained crosslinked and showed no relative sliding.

To gain further insights into motor behaviour at parallel or anti-parallel filament overlaps, we performed a microfluidics assay, in which myosins were first bound to surface-anchored individual filaments in the absence of ATP and ADP (Fig 3A). Second, we flowed in pre-formed short actin filaments of a different colour, which were captured by the myosins decorating the anchored filaments. Last, we switched to a solution containing ATP, ADP, and myosin-X to keep supplying motors, and observed the relative motion of the short filaments based on their polarity (Fig. 3B). Anti-parallel filaments slided at a velocity that almost matches the velocity of individual myosins on parallel bundles in the aforementioned microfluidics assay (Fig. 3C). Parallel filaments were almost static, with a sliding velocity of ∼ 4 nm.s^-1^ towards the barbed end of the anchored filament, which is also the direction of the flow. This slow sliding could originate from the viscous drag force applied by the microfluidics flow on the mobile filaments. We probed this by inverting the configuration, using Capping Protein to attach filaments by their barbed end to the surface of the chamber. Mobile filaments forming parallel overlaps with the anchored filaments slided in the direction of the flow at a comparable low velocity (Supp. Fig. S12). This indicates that myosins allow external forces applied to only a subset of filaments of a parallel bundle to reorganize the relative positioning of the filaments in the bundle. Those forming anti-parallel overlaps slided in the upstream direction at a velocity almost unaffected by the viscous drag (Supp. Fig. S12).

**Fig. 3.**
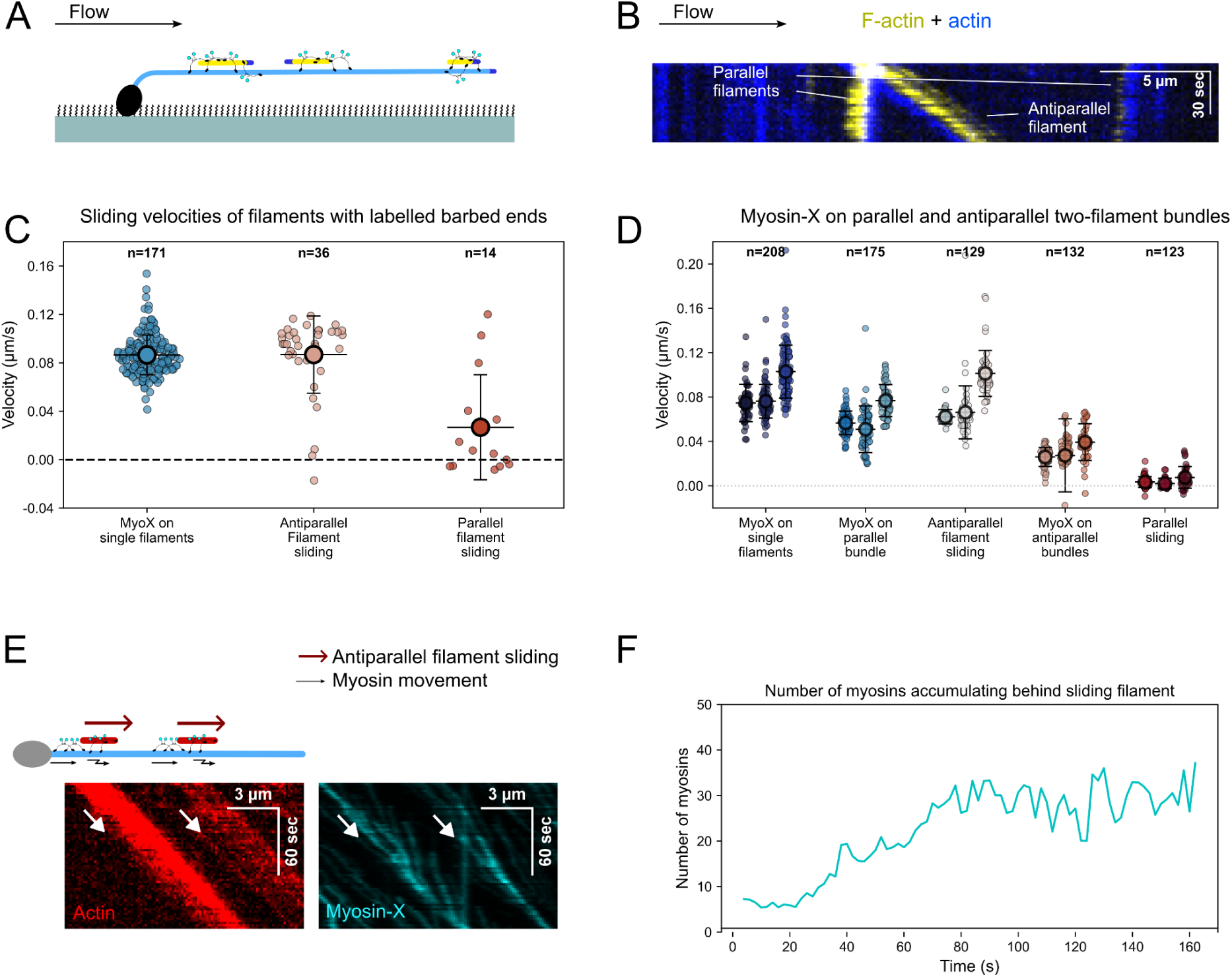
Myosin-X is processive on both parallel and anti-parallel 2-filament bundles. (A) Sketch describing the experiment where Alexa568-labeled actin filaments elongated from surface-anchored spectrin actin seeds in a microfluidics chamber are exposed to 75 nM myosin-X (10% myosin-X-mStayGold) and pre-polymerised Alexa647-labeled actin filaments, in the presence of 200 µM ATP and 800 µM ADP. Barbed ends are visualized by exposing them to 1 µM G-actin labeled in the same colour of the anchored filament, but in a higher labelling fraction, and 2 µM profilin, in the presence of 400 µM ADP for 2 minutes. Switching to the solution with myosin-X and ATP/ADP results in filament sliding. (B) Representative kymograph of an experiment described in A. (C) Myosin-X velocities on single filaments (n=171), and sliding velocities of filaments in anti-parallel (n=36) or parallel (n=14) configurations when forming bundles with anchored filaments. Large data points represent the average of each condition. Error bars are standard deviations. (D) Myosin-X velocities and sliding velocities of filaments in the different configurations. Velocity on antiparallel bundles is relative to the sliding filament. The sliding velocities relative to the anchored filament are shown in Supp. Fig. S13. Data from three independent experiments, at least 35 myosins were quantified in each condition for each experiment. Large data points represent the average of each condition. Error bars are standard deviations. (E) Kymograph showing the accumulation of myosin-X at the barbed ends of sliding filaments with unlabeled barbed ends (white arrows). (F) Accumulation of myosin-X over time near the barbed end of a sliding filament.

In the case of the anti-parallel configuration, in the reference frame of one of the filaments of the 2-filament bundle, myosin-X walks at half velocity usually observed on parallel bundles (Fig. 3D, Supp. Fig S12), indicating that motors have no preference for one filament and frequently change their walking direction. Moreover, as motors kept walking towards filament barbed ends, they accumulated near the barbed ends of the short sliding filaments (Fig. 3E). This makes sense because, as motors continue walking, they should reach slightly beyond the barbed end of the short filament, and, once there, as motors walk slightly faster on single filaments than anti-parallel filaments are sliding, motors should eventually re-enter the anti-parallel overlap region. Motors are thus ‘trapped’ at this boundary, while the short filament keeps sliding towards the barbed end of the anchored filament. Motor accumulation appears to saturate at around 26 (± 5) motors (n=5, Fig. 3F). While this number is comparable to the number of myosins in clusters observed in open chamber assays, we did not observe myosin cluster formation in this situation (at the motor concentration used). This is probably because motors are funneled from 2 filaments only which appears insufficient to increase motor density, and are not as packed as in larger bundles.

We assessed the stability over time of parallel and anti-parallel filament pairs, by growing individual pointed-end-anchored filaments with myosin-X in the absence of microfluidics. As individual filaments encounter each other with a random orientation (Fig. 4A), the surviving fraction of anti-parallel pairs dropped much more quickly than that of parallel pairs (Fig. 4B).

**Fig. 4.**
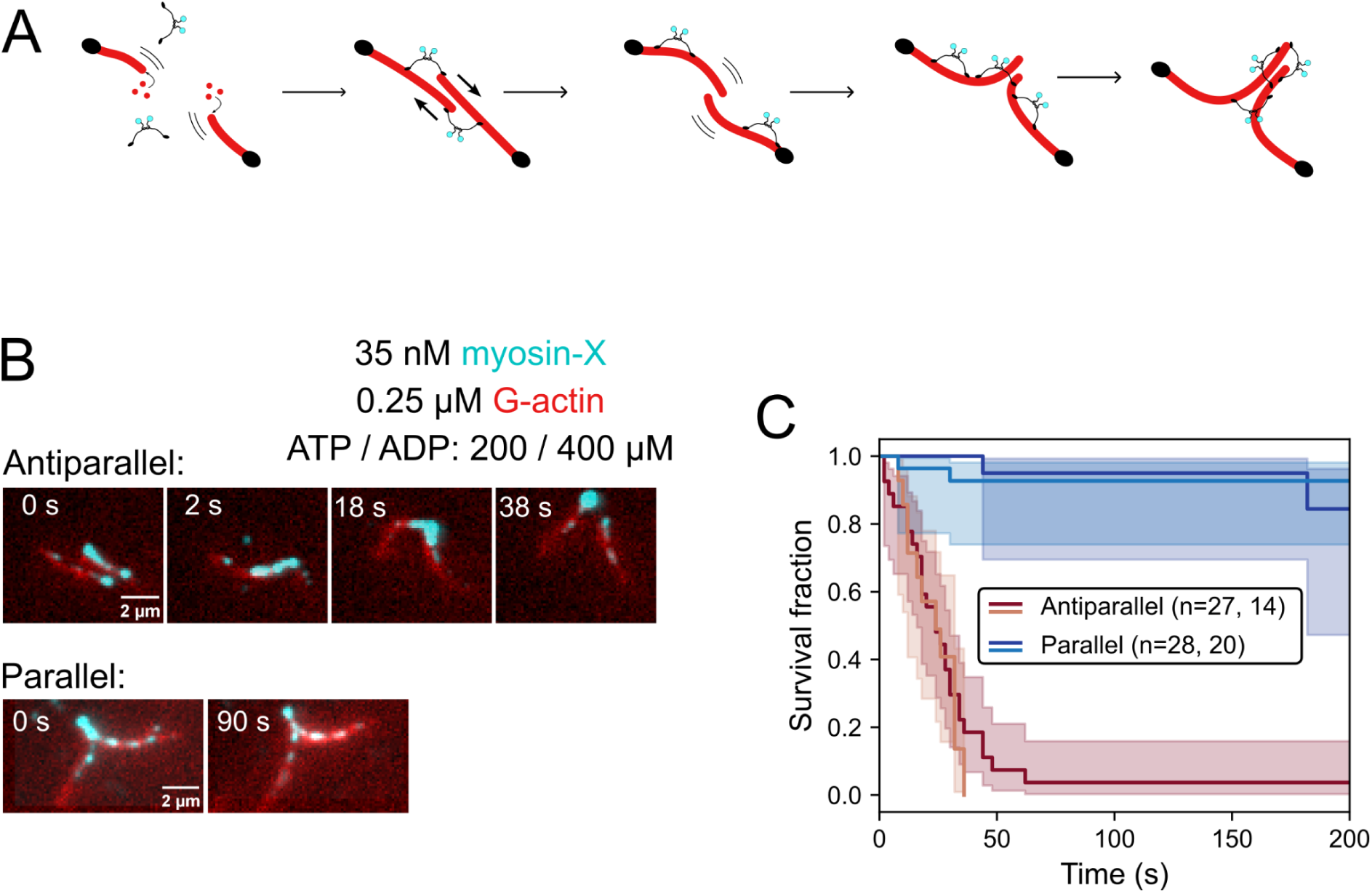
Myosin-X actively selects for parallel actin bundles. (A) Sketch describing the experiment to observe the outcome of parallel and antiparallel bundles from anchored filaments. Actin filaments were elongated from surface-anchored spectrin actin seeds (black circles) in an open-chamber assay, in the presence of myosin-X. (B) Snapshots from an experiment with 35 nM myosin-X, 0.25 µM Alexa568-labeled G-actin, in an F buffer containing 200 µM ATP and 400 µM ADP, and 0.2% methylcellulose. Scale Bar : 2 µm. Top: an antiparallel bundle forms and dissociates. Bottom: a parallel bundle remaining stable over time. (C) Survival fraction of parallel (n= 28 and 20) and anti-parallel (n=27 and 14) two-filament bundles as a function of time, from two independent experiments.

Overall, in the minimal case of 2-filament bundles mediated by myosin-X, myosin-X quickly sorts filaments according to their relative polarity.

### Density of myosin-X tunes actin filament elongation

We next asked whether myosin-X at bundle tips could affect barbed end dynamics. We performed microfluidics assays with single filaments in the presence of 2 µM ATP, to obtain a high density (∼ 4 /µm) of motors walking slowly (∼ 70 nm.s^-1^), so that motors are frequently near the barbed end of actin filaments. In such a condition, myosin-X reduced both barbed end depolymerization and elongation rates to the same extent, by ∼ 30 % (Fig. 5A, B).

**Fig. 5.**
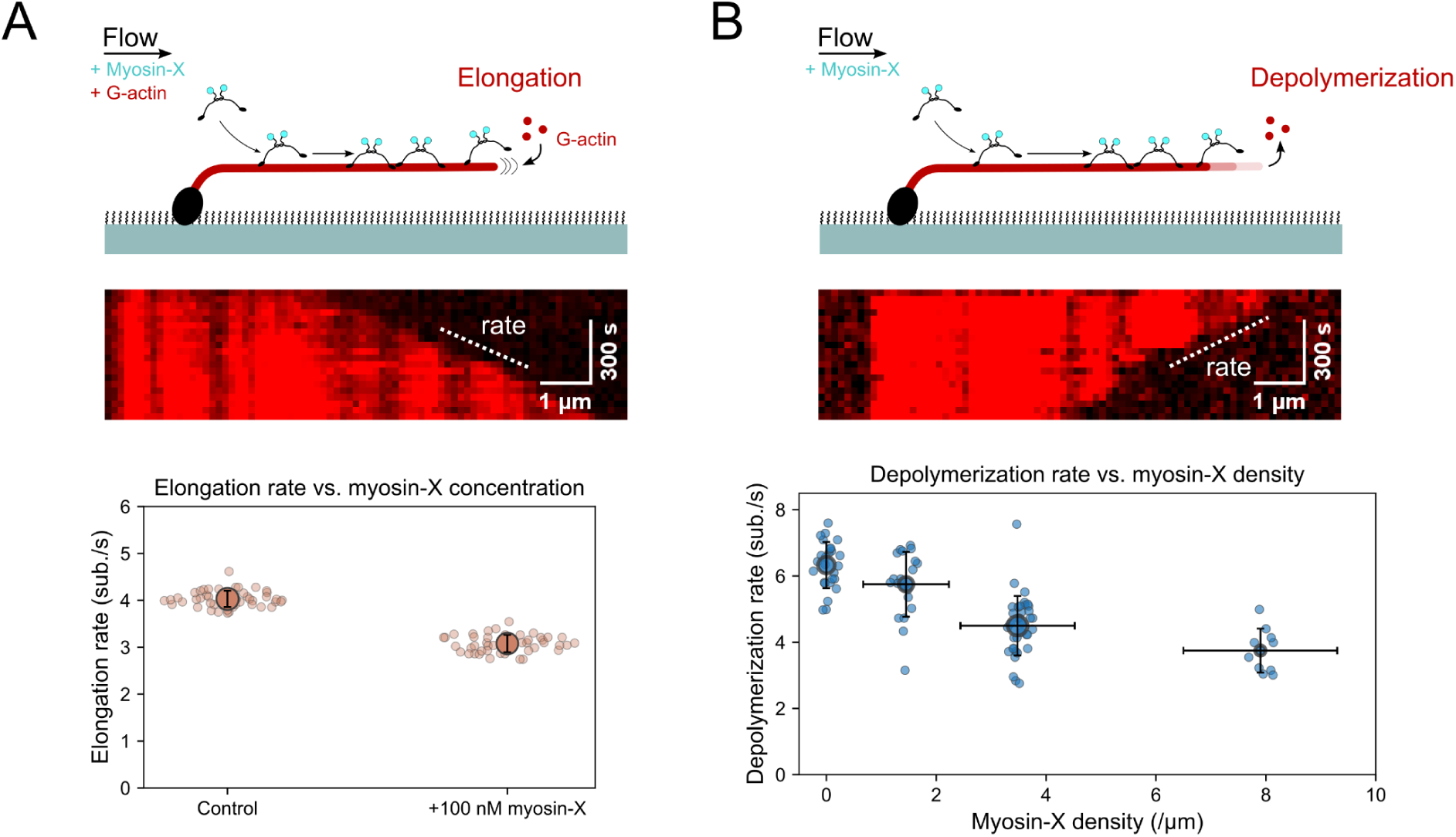
Myosin-X tunes barbed end dynamics of single actin filaments. (A) Barbed end elongation rate in the presence of 0.5 µM actin (n = 44), and with the addition of 100 nM myosin-X (n= 44), with 2 µM ATP in solution, measured from single actin filaments elongating from surface-anchored spectrin-actin seeds in a microfluidics assay. Error bars are standard deviations. (B) Barbed end depolymerization rate in the absence of actin (n= 26), and with increasing concentrations of myosin-X (50 (n=20), 100 (n=36), and 200 (n=10) nM, respectively), with 2 µM ATP in solution, measured in a similar experiment to (A). Myosin-X density is determined by measuring the fluorescence intensity along filaments. Error bars are standard deviations.

Myosin-X affected barbed end dynamics in a concentration-dependent manner (Fig. 5B). Furthermore, myosin-X totally abolished barbed end elongation in the absence of ATP (Supp. Fig. S14).

Assuming that myosin-X can bind equally well to actin subunits near or at the barbed end as to subunits along the filament, the direct landing of myosin-X near the barbed end accounts for ∼ 1/6 of the myosins reaching this region, the rest are the ones that land upstream, and reach the barbed end by walking processively. Neglecting direct targeting, and considering only that the filament collects and delivers myosins to the barbed end, the probability of the barbed end being blocked by a walking motor is τ*J, where τ is the residence time of a motor near the barbed end blocking its elongation/depolymerization, and J is the motor flux (Supp. Fig. S14). Therefore, a reduction of the elongation rate by ∼ 30 % gives an estimate of the motor residency duration τ = 0.7/J ∼ 1.5 seconds. Considering the velocity of myosin-X (70 nm.s^-1^) and its maximum span of the fully extended motor lever arm (∼ 50 nm), this value indicates that motors could slow down or pause at the barbed end.

## Discussion

Myosin motors can transform passively cross-linked actin architectures, shaping actin networks by applying mechanical tension leading to filament reorganisation (Murrell et al., 2015). The most well-known illustration is the role of myosin-II mini-filaments in the actin cortex (Cheffings et al., 2016; Lehtimäki et al., 2021; Truong Quang et al., 2021; Vignaud et al., 2021).

Myosin-X is a dimeric, high-duty-ratio motor that can take steps of various sizes, depending on the organization of the actin filaments (Berg et al., 2000; Homma and Ikebe, 2005; Ropars et al., 2016; Sun et al., 2010; Tokuo and Ikebe, 2004). Here, we show that the motor activity of dimeric myosin-X, even in the absence of its cargo-binding tail, combines processive movement with the sorting and bundling of actin filaments into parallel arrays with gathered barbed ends (Fig. 6A). The funneling of motors towards bundle tips increases the local motor density, tunes barbed end dynamics, and may lead to the dynamic clustering of myosins (Fig. 6B). Together, these properties provide a molecular basis for the initiation of filopodia by myosin-X motors (Tokuo et al., 2007), and for the large accumulation of myosin-X at filopodia tips (Berg et al., 2000; Berg and Cheney, 2002; Tokuo and Ikebe, 2004).

**Fig. 6.**
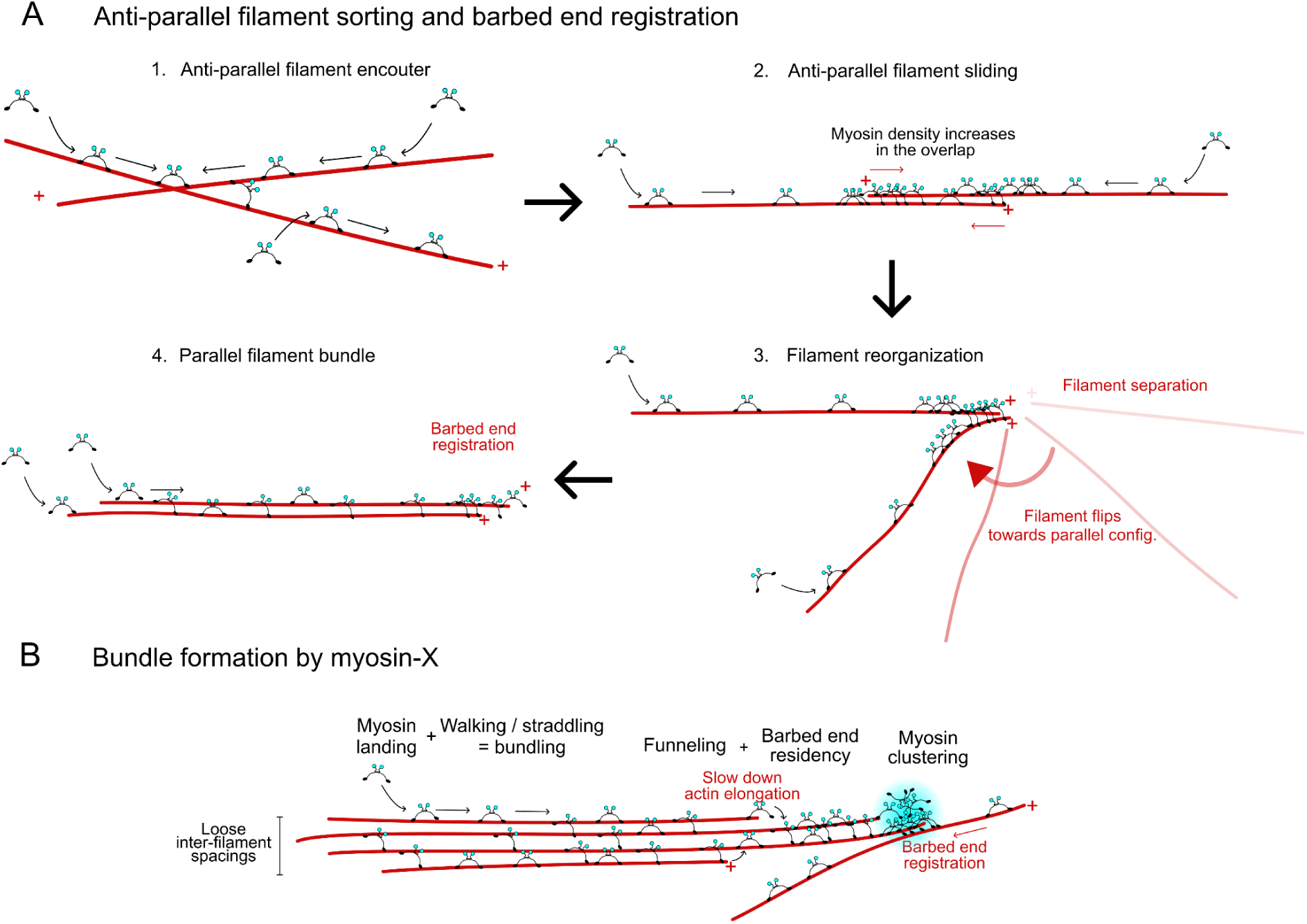
Initiation of parallel actin bundles by Myosin-X. (A) Sequence of events leading to parallel bundle formation: 1. Two freely diffusing filaments are crosslinked by myosin-X in an anti-parallel configuration; 2. Myosin-X slides filaments apart and accumulates in the overlap region, in particular near barbed ends of both filaments; 3. After complete sliding, filaments separate with their barbed end in close proximity. Accumulated myosin-X allows filaments reorientation while maintaining barbed ends in close proximity; 4. The parallel configuration is stable and barbed ends are in register. (B) The different features leading to large parallel actin bundle formation by myosin-X: 1. myosin-X efficiently straddles along two filaments of a bundle and processively walks towards their barbed ends; 2. Funneling of myosin-X leads to the increase of motor density towards the bundle tip, and slows down filament elongation; 3. Myosin-X clusters above a critical motor density; 4. A new filament can be incorporated in the bundle, with its barbed end registered, as the myosin cluster at the bundle tip pulls the filament backward.

Bundle formation is dependent on myosin, ATP, and ADP concentrations. This ability likely arises from the capacity of myosin-X to side-step, as reported by many studies on fascin-actin bundles (Nguyen et al., 2023; Ropars et al., 2016; Sato et al., 2017; Sun et al., 2010), and from its high duty ratio (Homma and Ikebe, 2005), since increasing ADP concentration raises the duty ratio and the probability that both motor domains are bound to actin subunits simultaneously. Furthermore, increasing motor density on actin filaments could lead to a feed-forward mechanism, in which more motors cross-linking filaments increase the probability of side-stepping of other motors, further reinforcing filament crosslinking.

To form parallel actin bundles with gathered barbed ends, myosin-X needs to perform a few operations based on the polarity of actin filaments.

First, the extended lever arm of myosin-X seems to provide enough flexibility for the motor to crosslink two filaments regardless of their relative orientation. Myosin-X crosslinks anti-parallel filaments while remaining processive, so that its power-stroke efficiently slides filaments of opposite polarities apart. This expected behaviour could lead to bundle contractility or extension depending on the exact bundle geometry, as observed experimentally for myosin-II thick filaments (Murrell and Gardel, 2012; Thoresen et al., 2011) and described theoretically (Lenz et al., 2012; Zemel and Mogilner, 2009). As anti-parallel filaments slide apart, myosin-X accumulates in the shrinking overlap region, in the vicinity of the barbed end of both filaments. Such local accumulation sustains persistent sliding. This could also seed a high motor density at barbed ends, which would later be funneled towards the tip of a nascent bundle. Above a critical density threshold, traffic jams or motor clustering may appear.

How can barbed end gathering of filaments in parallel bundles be achievedIn the case where anti-parallel filaments have almost slid apart entirely, we believe that accumulated motors at barbed ends could serve as a strong link between filament barbed ends, so that filaments remain connected while they can pivot to re-orient in the parallel configuration (Fig. 6A). This behaviour is reminiscent of single non-muscle myosin-II mini-filaments forming bundles or asters in vitro (Wollrab et al., 2018), where the high number of motor domains held together in a mini-filament would play the role of myosin-X clustering reported here.

Second, Myosin-X maintains a loose and dynamic connection between parallel filaments, which accommodates filament sliding if an external force is applied, as revealed in our assay by the flow-driven sliding of mobile filaments along anchored filaments. Though, the behaviour of myosin-X differs markedly from that of passive crosslinkers, such as anillin for actin filaments, or Ase-1 for microtubules, which expand filament overlap by an entropy-driven mechanism (Kučera et al., 2021; Lansky et al., 2015).

Here, in contrast to previous studies on myosin-X with various dimerization strategies (Nagy et al., 2008; Nguyen et al., 2023; Ropars et al., 2016), our myosin-X construct displays only limited differences in terms of velocity and run length on single filaments, on the ‘loose’ bundles it creates (both parallel and anti-parallel), and on more rigid fascin bundles. Myosin-X is thus not specifically tailored to walk on fascin-actin bundles. It accommodates a broad range of inter-filament spacings, including those it generates.

The increased density of myosin-X at bundle tips has a direct impact on actin assembly dynamics. At the single filament level, myosin-X slows down both barbed end elongation and depolymerization in a motor density-dependent manner. Our estimate of the residence time of motors in the vicinity of the barbed end (τ ∼ 1.8 seconds; for a motor velocity along filaments of ∼ 70 nm.s^-1^) indicates that motors probably slow down or pause at the barbed end. To our knowledge, this is the first report, though indirect, of the residency of a dimerised myosin motor on actin filament ends. As myosin-X approaches the barbed end and the ‘standard’ 36 nm step size is no longer available, the motor may be able to take smaller steps and continue walking, or even to perform backwards steps to remain bound to the filament for a longer time, before finally detaching. In such a proximity with the barbed end, the motor would sterically interfere with subunit addition and removal. This activity is in contrast to the depolymerase activity of monomeric myosin-Ib observed in gliding assays (Pernier et al., 2019).

The regulation of actin assembly dynamics by myosin-X is reminiscent of, but yet distinct from, that of myosin-XV in the strongly bound state (i.e. apo or ADP-bound state) where it increases filament nucleation but slows down polymerization (Gong et al., 2022; Moreland et al., 2025). Myosin-X completely abolishes barbed end elongation in the strongly bound state, a stronger effect than that of myosin-XV. Besides, we show that the reduction of barbed end dynamics can be achieved by processive myosin-X reaching the barbed end, in the presence of ATP and absence of ADP.

The funneling of motors towards remaining filaments at bundle tips efficiently increases motor density towards bundle tips. This thus provides a way to increase the net flux of motors per filament towards the barbed end, and could potentially lead to a jamming situation because of the myosin-X increased residency at the barbed end. Above a certain density, motors form clusters that remain cohesive even after detaching from bundles, and that only partially exchange with the pool of soluble motors. In cells, this clustering may be further amplified. Indeed, the close configuration of filopodia tips and the membrane attachment of myosin-X via its tail PH domain, would confine motors and slow down their diffusion. This could allow more efficient re-binding of the motors that unbind from bundle tips, and thus maintain a high local motor density. It is tempting to relate the dynamic clusters we observe in vitro to the myosin-X-rich puncta observed at filopodia initiation sites and filopodia tips, although myosin-X’s binding partner Ena/VASP plays an important role in effective filopodia maturation (Bohil et al., 2006; Pokrant et al., 2023; Tokuo et al., 2007).

Since myosin-X follows a helical path around actin filaments (Arsenault et al., 2009; Sun et al., 2010), the twirling of filaments within myosin-X-actin bundles could, in principle, generate torsional stresses, potentially inducing filament breakage and subsequent rearrangements. In our assays with free filaments, bundles sometimes looked curly (Supp. Movie 3) which could indeed indicate such twirling motion. This observation is in line with the cork-screw shape of retracting filopodia in cells, attributed in part to myosin-X activity (Leijnse et al., 2022, 2015).

Together, our results support a model in which the motor activity of myosin-X is sufficient to initiate the formation of parallel actin bundles. Myosin-X sorts filaments based on their polarity, gathers their barbed ends, and concentrates at bundle tips, where it tunes barbed end dynamics. In cells, this activity would precede the recruitment of fascin, which would subsequently consolidate the nascent bundle into a rigid, filopodia-competent architecture. The gathering of barbed ends and the local clustering of motors could, in turn, provide a platform for an efficient recruitment of elongators such as Ena/VASP at filopodia initiation sites (Jacquemet et al., 2019; Miihkinen et al., 2021; Pokrant et al., 2023; Popović et al., 2023). Our assays rely on a tailless, constitutively dimerized construct. How full-length myosin-X, whose dimerization at the membrane and cargo binding are regulated in cells, performs these operations remains to be explored. How the interplay between myosin-X, passive crosslinkers, and elongators, such as Ena/VASP, orchestrates the transition from bundle initiation to filopodium extension will be an exciting direction for future work.

## Supplementary Figures

**Supp. Fig. S1.**
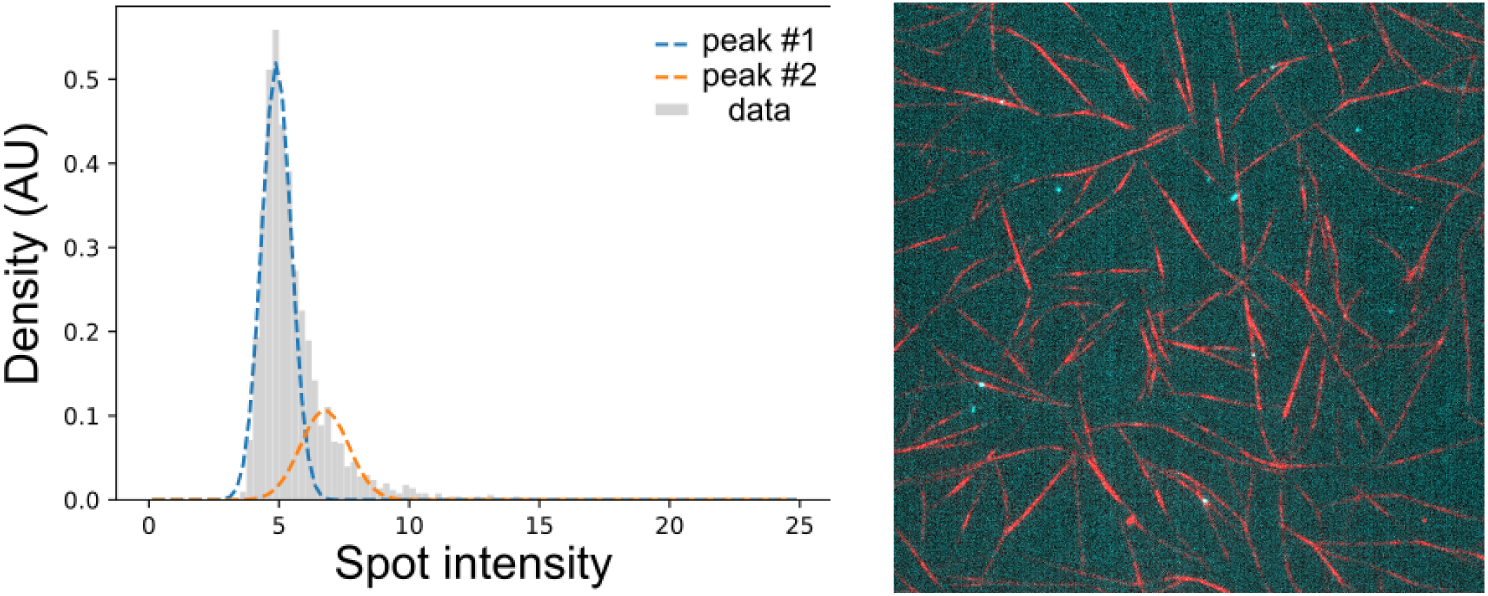
Fluorescence of myosin-X spots on actin bundles. Distribution of myosin-X spot intensities, for 2026 spots automatically detected by the TrackMate/FiJi plugin, from images acquired in an open-chamber assay with 5 nM myosin-X processively walking on fascin-actin bundles. The distribution is best fitted with two Gaussian distributions with a relative weight of 75% and 25%.

**Supp. Fig. S2.**
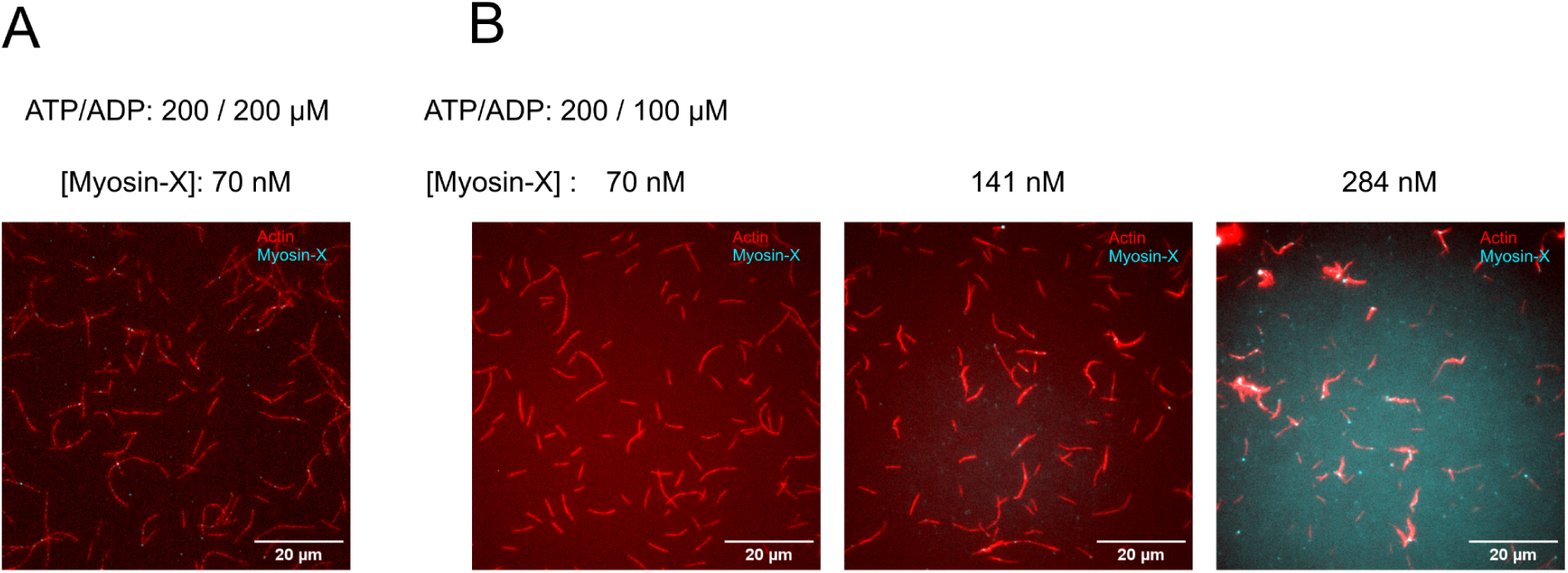
Bundling by myosin-X depends on the motor concentration. (A) At 200 / 200 µM ATP/ADP, 70 nM myosin-X is not sufficient to induce bundling. (B) At 200 / 100 µM ATP/ADP, a high concentration of myosin-X can induce bundling.

**Supp. Fig. S3.**
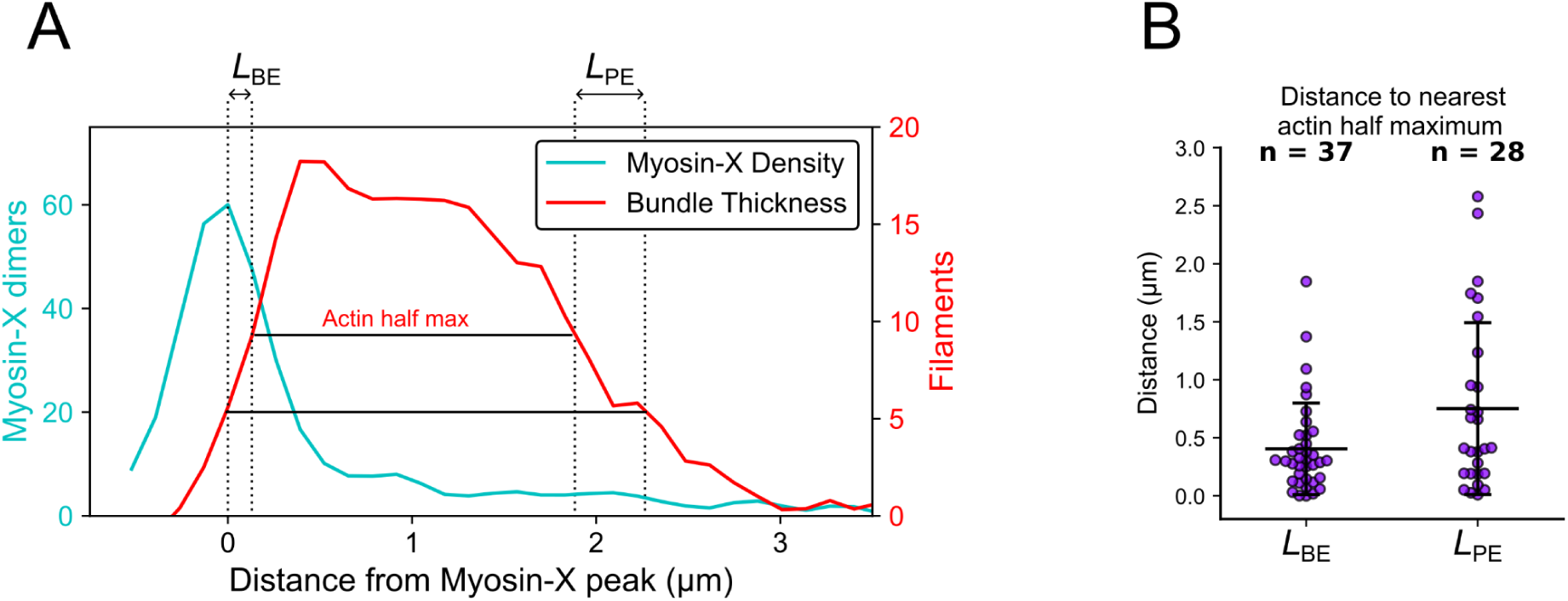
Barbed end registration in bundles organised by myosin-X. (A) Typical intensity profiles of myosin-X and actin along a bundle, normalised to individual myosin-X dimers and actin filaments. *L*_BE_ is defined as the distance between the highest point of the myosin-X peak to the first actin half maximum. *L*_PE_ is defined as the distance between the actin half maximum on the other side of the bundle and the point where the actin intensity is equal to that at the myosin-X peak. (B) Distributions of *L*_BE_ and *L*_PE_. *L*_BE_ is smaller than *L*_PE_, indicating that barbed ends are registered. Data from two independent experiments (same as Fig. 1D). Error bars are standard deviations.

**Supp. Fig. S4.**
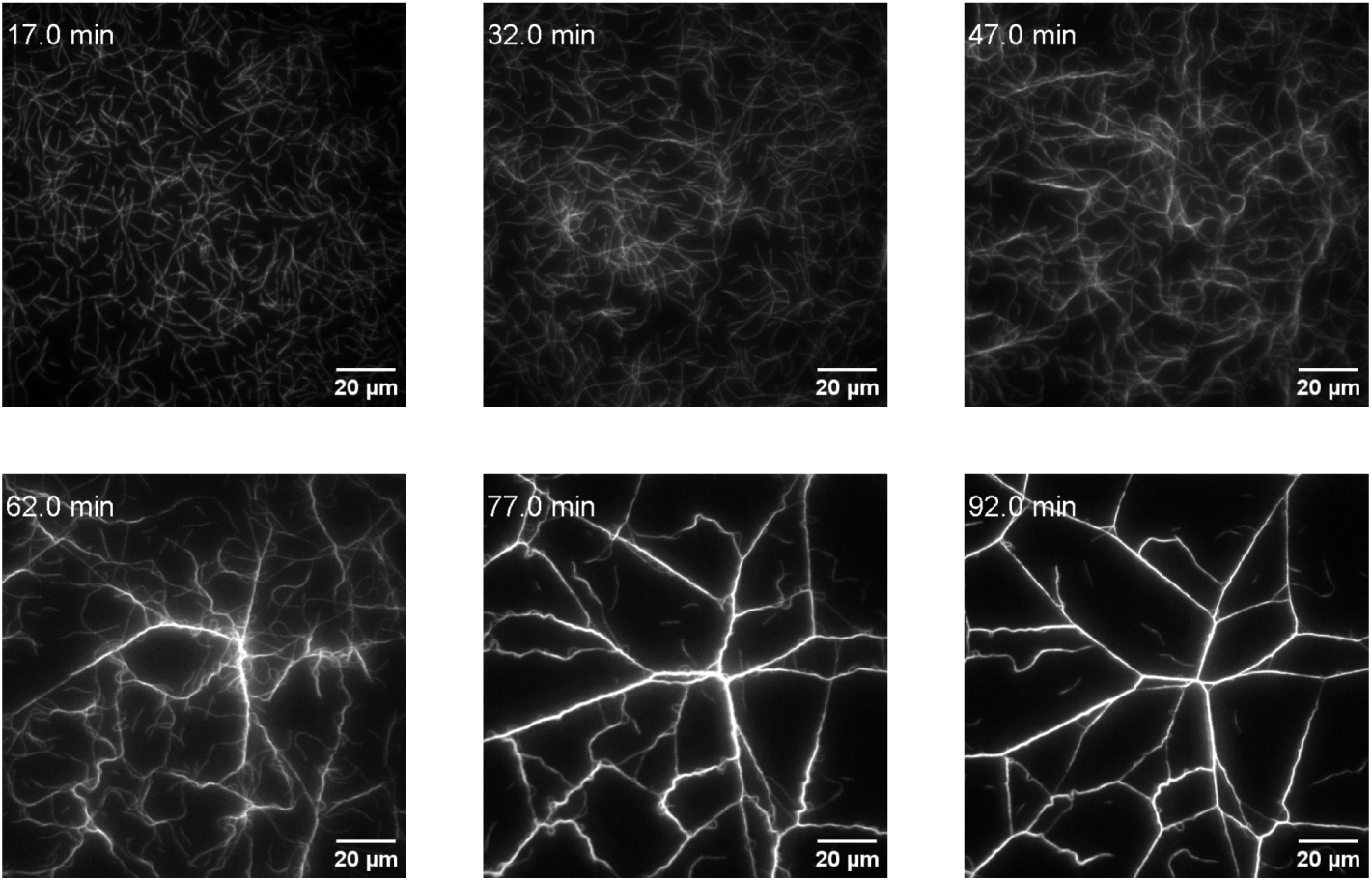
**Formation of contractile actin networks by myosin-X.** A high density of long actin filaments (∼14 µm) and the presence of 60 nM unlabeled myosin-X, with a buffer containing 20 µM ATP (no regeneration system), leads to the formation of contractile actin networks. Contrast settings are identical in all images. Scale bar: 20 µm.

**Supp. Fig. S5.**
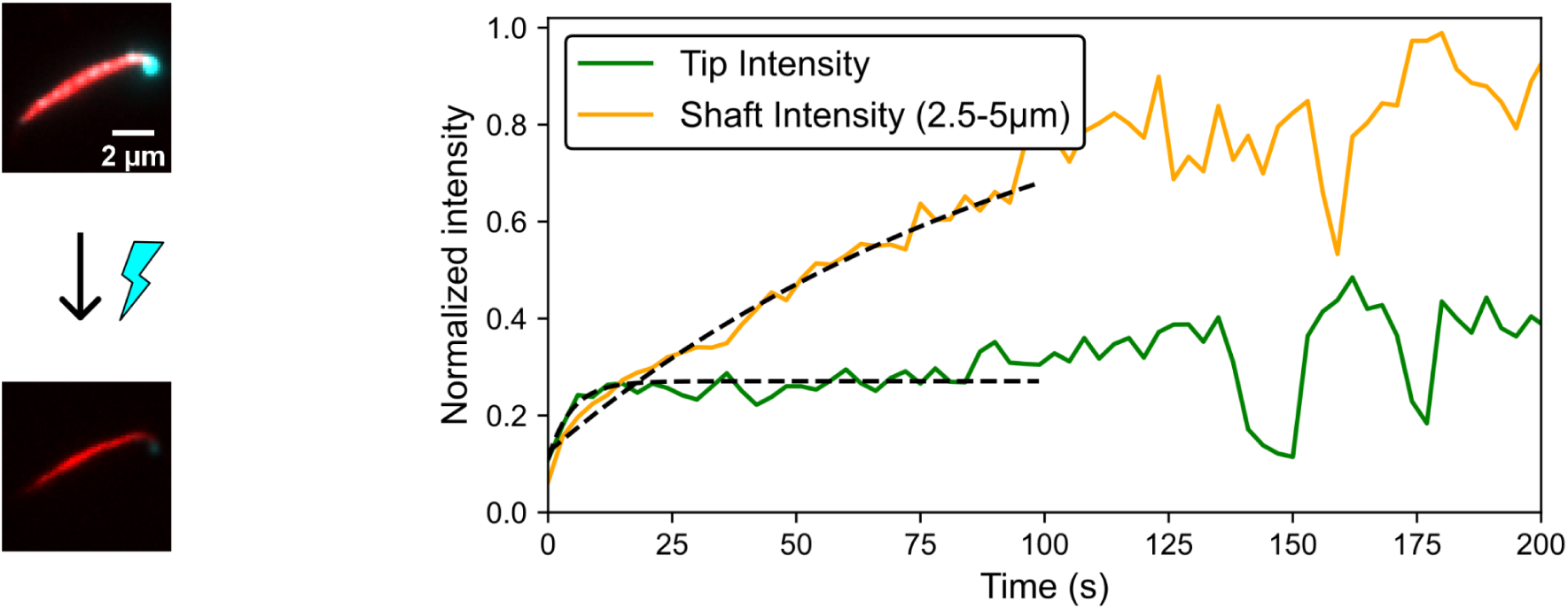
Fluorescence recovery of myosin-X at the tip and along the shaft of a bundle. FRAP on a complete myosin-X-induced actin bundle, not adhered to the glass surface. Shown are the myosin intensities normalised to the average intensity before photobleaching. Myosin-X fluorescence intensity was integrated across the first 2 µm from the bundle tip. The myosin-X fluorescence intensity along the shaft is the average intensity integrated along the bundle 2.5 to 5 µm away from the tip.

**Supp. Fig. S6.**
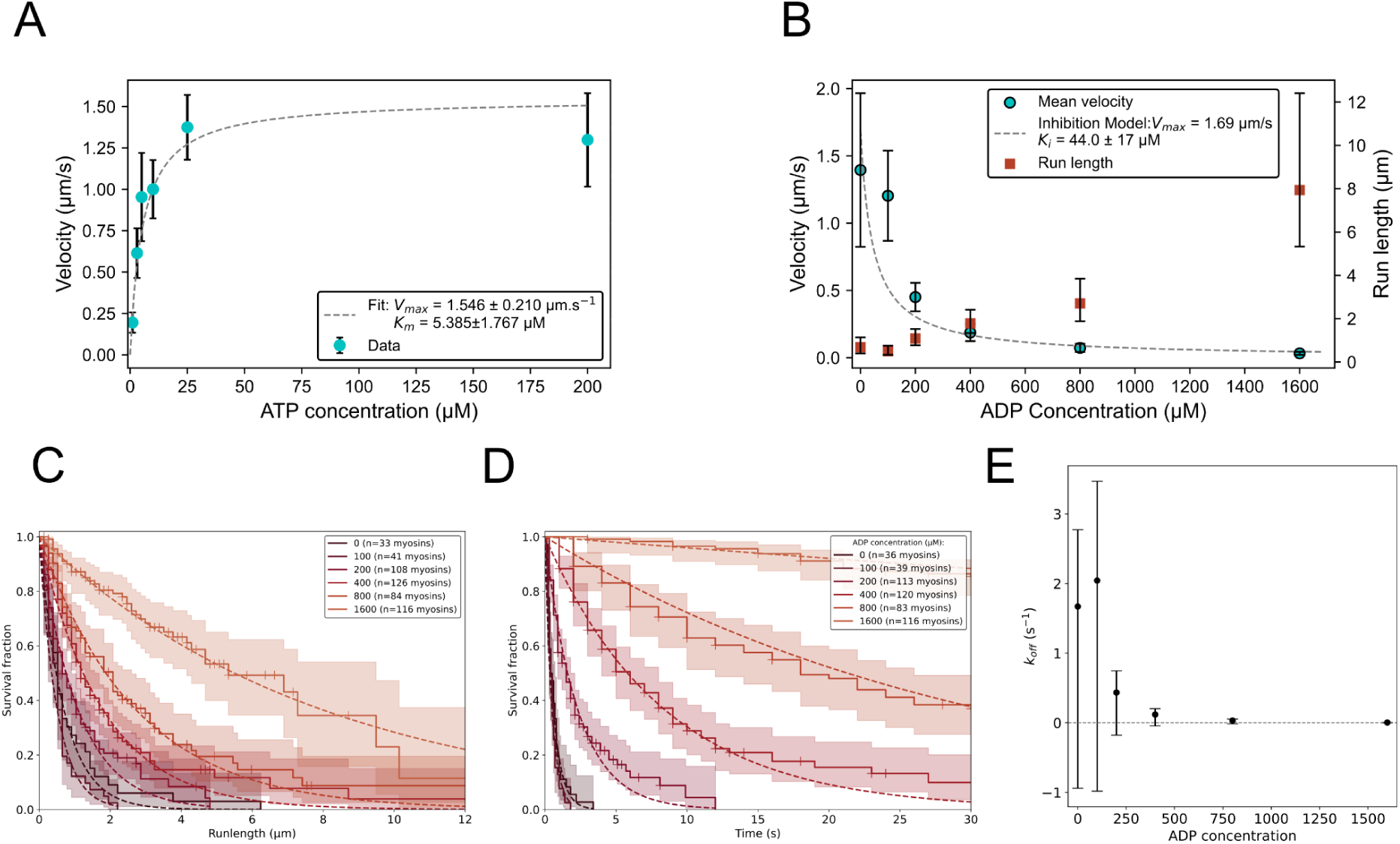
Biochemical characterisation of myosin-X-mStayGold construct. (A) Velocity as a function of ATP concentration. Experiments were performed in microfluidics on single filaments, in F-buffer, with an ATP regeneration system and the desired Mg-ATP concentration. Error bars are standard deviations. Fit with a simple Michaelis-Menten equation leads to the estimation of the apparent affinity *K*_m_ = 5.4 (± 1.8) µM and V_max_ = 1.5 µm/s. (B) Velocity and run length as a function of ADP concentrations. All experiments were performed in F-buffer, with 200 µM Mg-ATP, and different concentrations of Mg-ADP. Run lengths were determined from Kaplan-Meier fits from survival curves. Error bars for the velocity are standard deviations. Error bars for the run lengths are 95% confidence intervals. (C) Kaplan-Meier survival curves of myosin-X run length at different ADP concentrations in the presence of 200 µM ATP, resulting in the characteristic run lengths shown in B. (D) Survival fractions of myosin-X processivity at different ADP concentrations, at 200 µM ATP. (E) *k*_off_ values of myosin-X at different concentrations of ADP at 200 µM ADP, from the survival curves in D. Error bars are 95% confidence intervals.

**Supp. Fig. S7.**
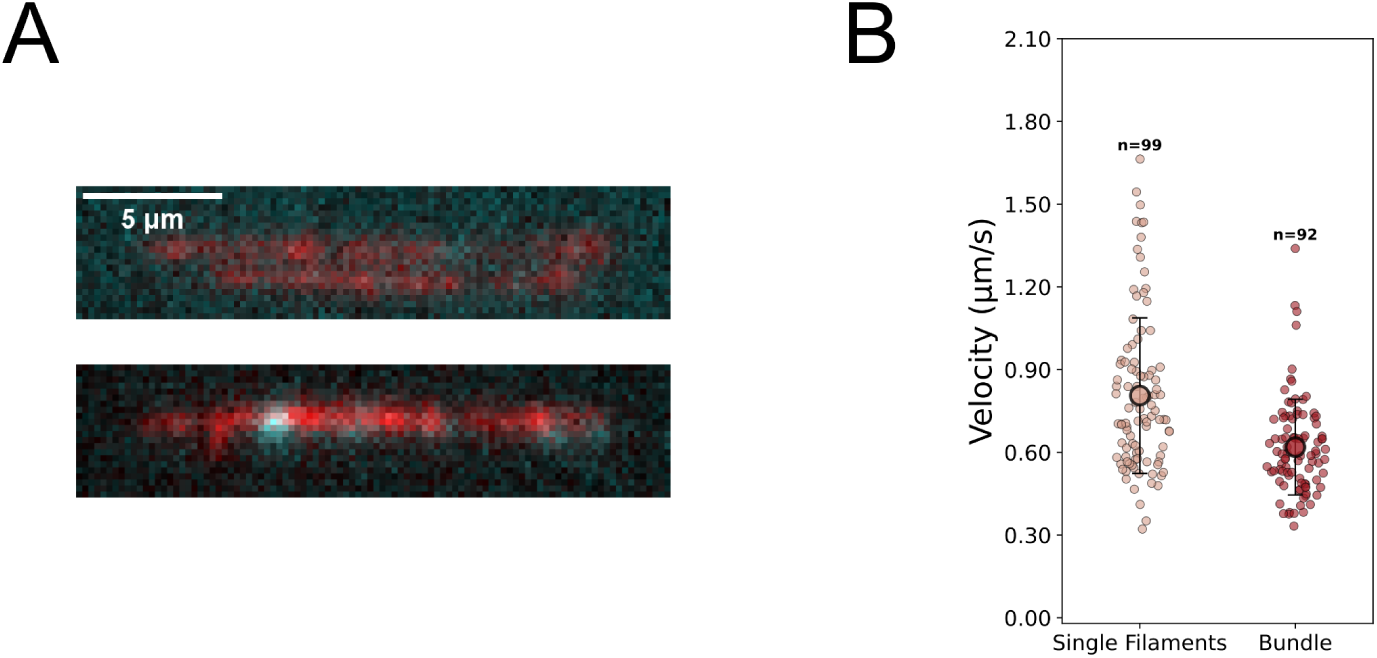
Myosin-X induces bundles in microfluidics. (A) In a microfluidics assay, actin filaments elongated from surface-anchored spectrin-actin seeds are exposed to 0.5 µM myosin-X in a buffer containing 200 µM ATP (without ATP regeneration system). (B) Velocity of myosin-X on single filaments (n=99) and 2-filament actin bundles induced by myosin-X (n=92). The velocity on bundles is 17% lower than on single filaments. Large dots represent averages and error bars are standard deviations.

**Supp. Fig. S8.**
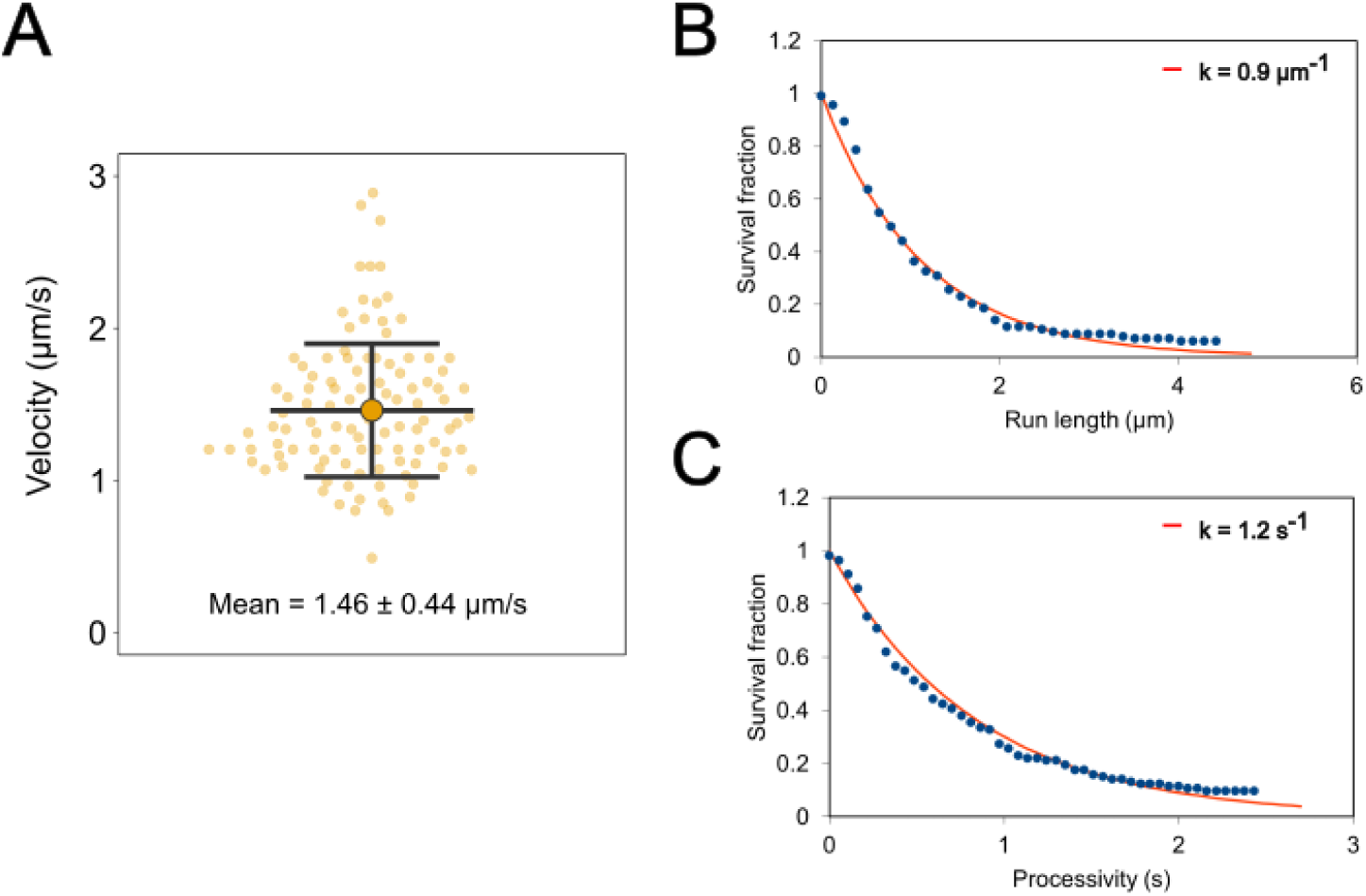
Walking behaviour of myosin-X on large fascin-actin bundles in open chambers, at 200 µM ATP. (A) Velocity of individual myosin-X (n = 113 myosins). The large data is the average of the population. Error bars represent standard deviations. (B and C) Survival curves of run length (n = 106 events) and processivity (n = 102 events) fitted by single exponential decay functions.

**Supp. Fig. S9.**
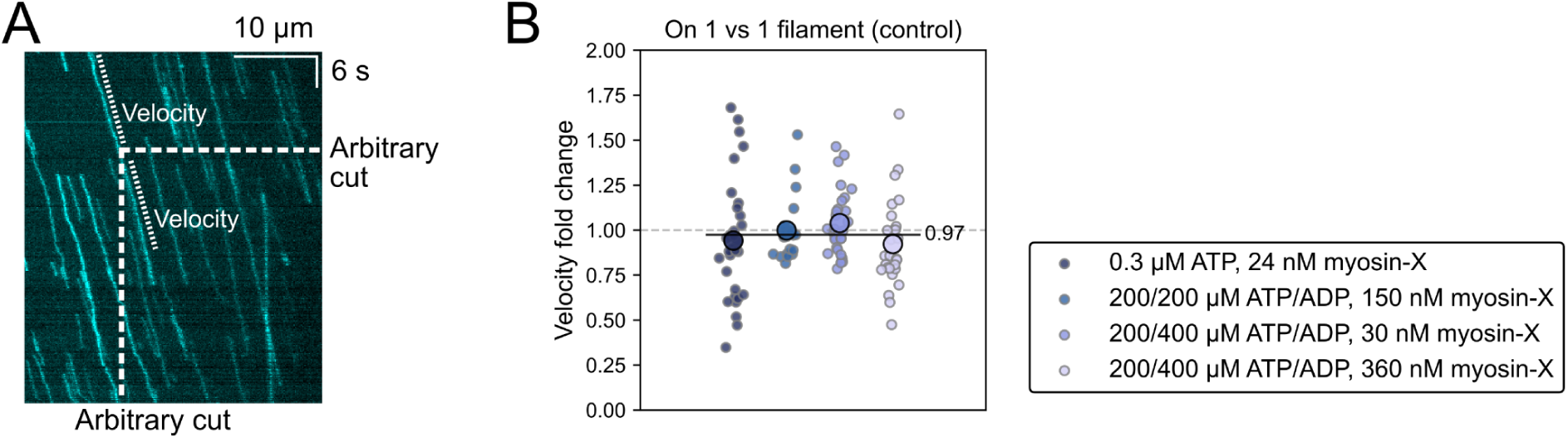
Analysis control: ‘switching’ between single filaments. (A) Kymograph of myosin-X walking on a single filament in microfluidics (actin not shown). As a control to the analysis presented in Fig. 2B, myosin tracks on single filaments were cut in two at an arbitrary position and both parts were analysed, and the fold change was calculated. (B) From left to right: n = 66, 44, 58, 54 myosins, from the same experiments as Fig. 2B. Calculated fold changes in myosin-X velocities before and after arbitrary boundaries in kymographs from single filaments. The ratio was calculated as in Fig. 2B. Noise indicates error in manual measurements + fluctuations in myosin velocity.

**Supp. Fig. S10.**
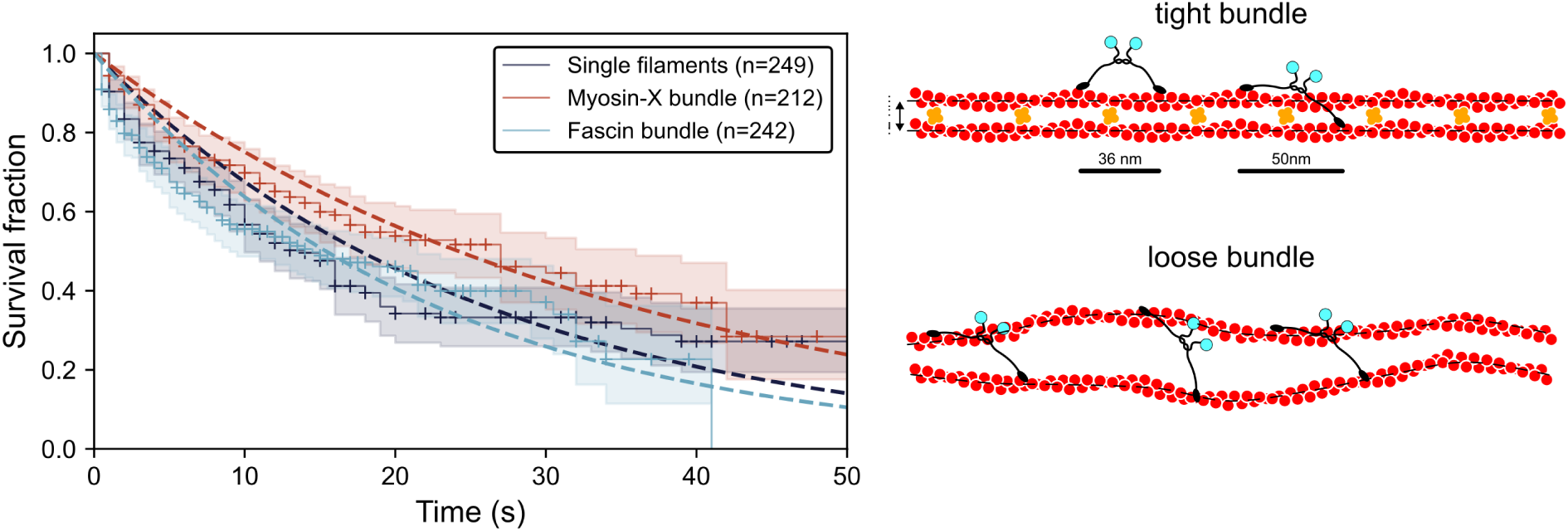
Processivity of myosin-X on single filaments, myosin-X bundles and fascin bundles, in the presence of 200 µM ATP and 400 µM ADP. (left) Dwell time of individual myosin-X on single filaments: 25.5 s (95% CI: 18.7 - 34.3 s), on myosin-X bundles: 34.9 s (95% CI: 25.9 - 47.7 s), and on fascin bundles: 22.2 s (95% CI: 17.0 - 28.8 s). (right) Schematics illustrating the difference in inter-filament spacing between bundles induced by fascin or by myosin-X.

**Supp. Fig. S11.**
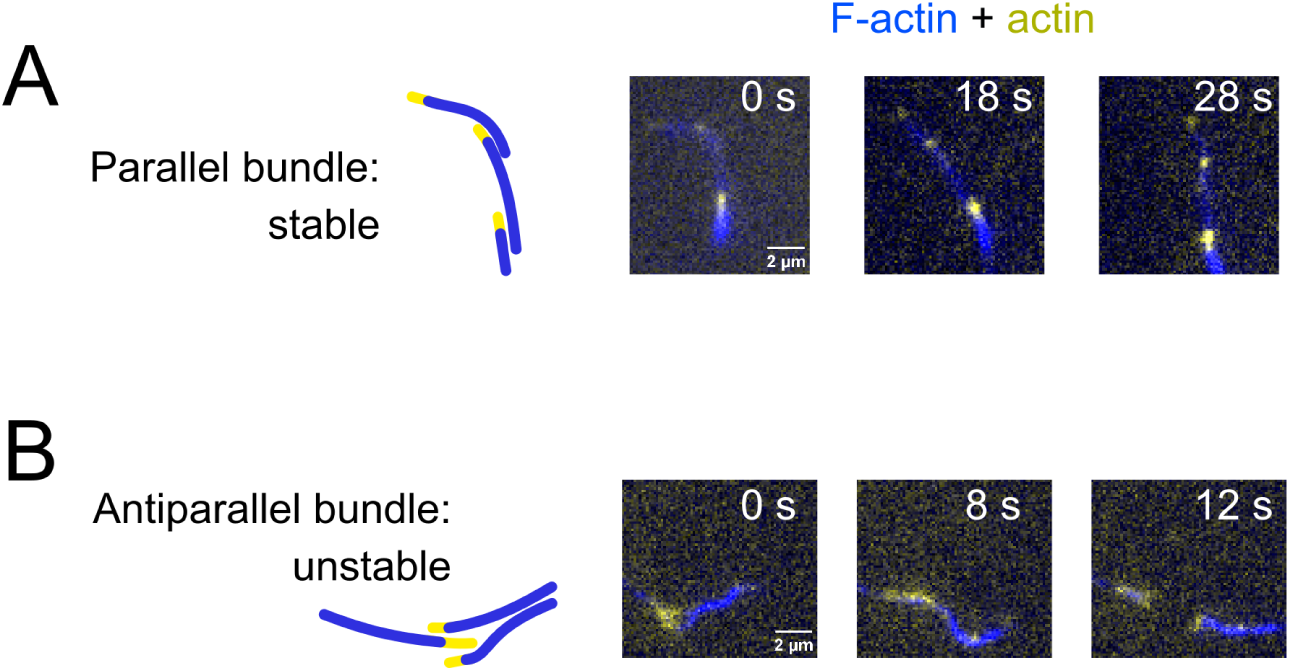
Sliding filaments in open chambers. Actin-Atto643 was polymerised at 20 µM for 20 min, then diluted to 0.5 µM in presence of an additional 0.25 µM G-actin-Atto488 to label barbed ends. 7.5 nM filaments were injected into an open chamber. 150 nM G-actin-Atto488 and 75 nM unlabeled myosin-X were added in a second injection in the presence of 200 µM ATP and 400 µM ADP.

**Supp. Fig. S12.**
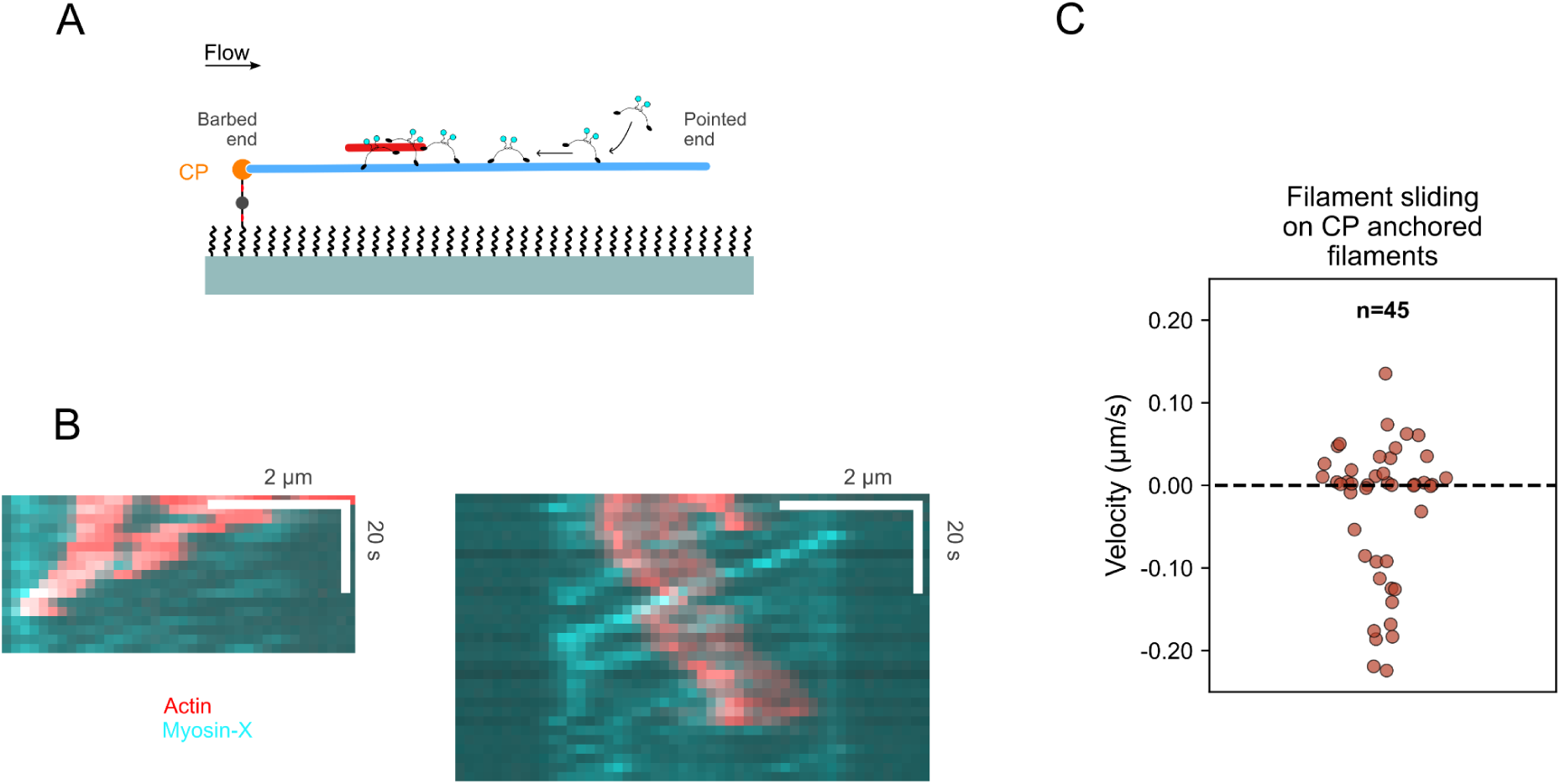
Sliding velocities of pre-polymerized filaments exposed to Capping Protein-anchored filaments decorated with myosin-X. (A) Experiment in microfluidics similar to Fig. 3. Filaments were anchored to the surface with Capping Protein. In this setup, myosin-X walks against the flow. (B) Kymographs showing filaments sliding upstream (leftwards) and downstream. The anchored filament is not shown. (C) Filament sliding velocities in the presence of 60 nM myosin-X-mStayGold, at 200 µM ATP and 400 µM ADP. Negative velocity indicated sliding against the flow, towards the barbed end of the anchored filament. The distribution is two-sided, suggesting that antiparallel sliding is persistent even when against the flow. Filaments sliding slowly in the direction of the pointed end of the anchored filaments are expected to be parallel. When filaments are anchored at their pointed ends, parallel pre-polymerised filaments also slide in the direction of the flow (Fig. 3C).

**Supp. Fig. S13.**
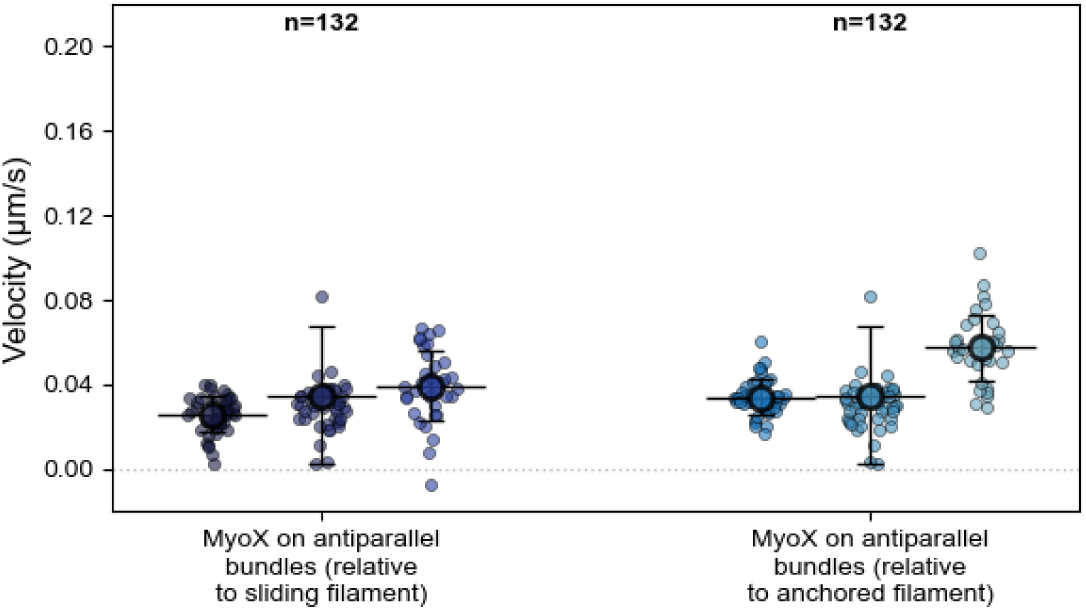
Myosin-X walking velocity on antiparallel bundles, relative to the sliding filament or the anchored filament. Data from the same three independent experiments as in Fig. 3D.

**Supp. Fig. S14.**
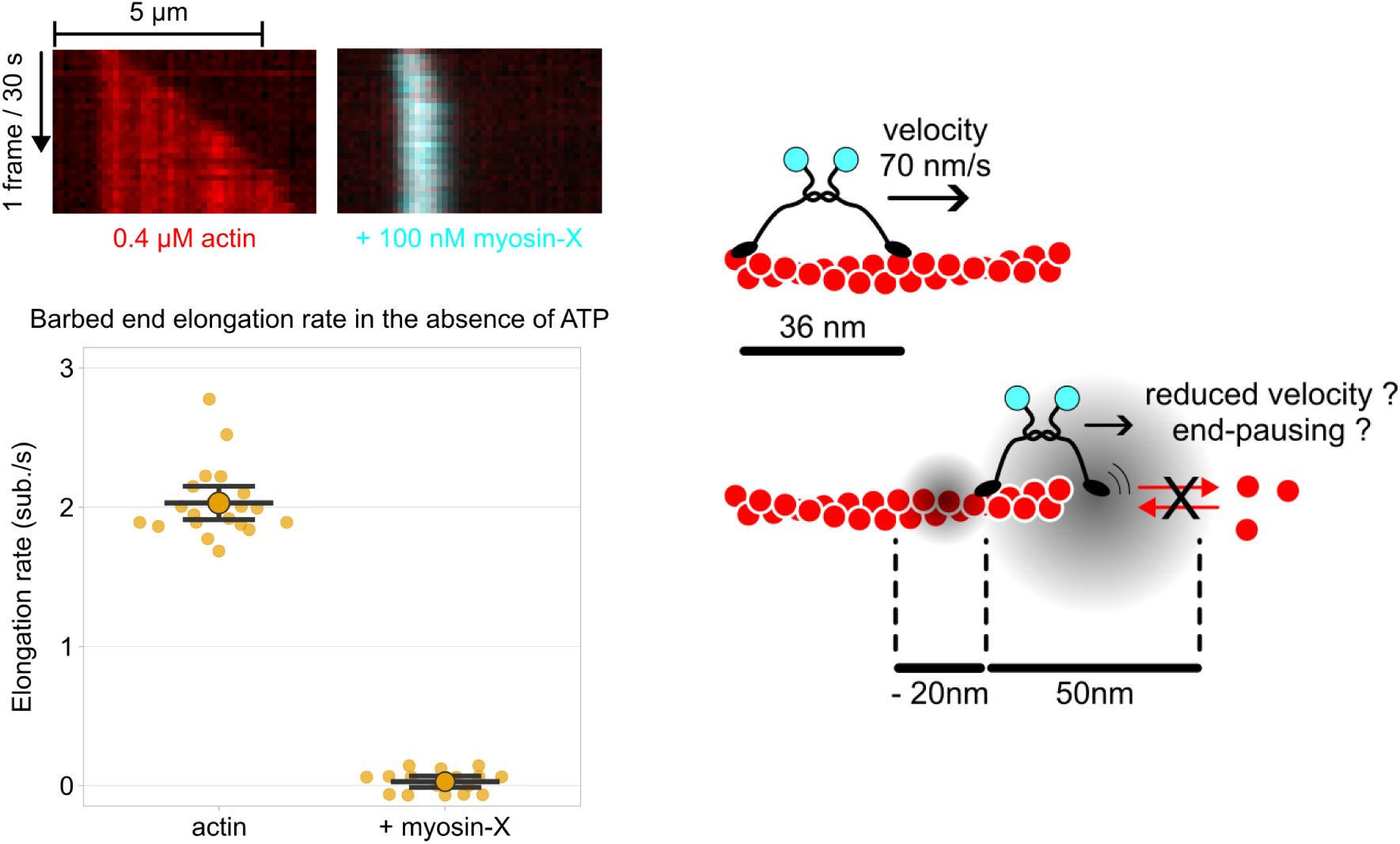
Myosin-X blocks barbed end elongation in absence of ATP. (left) In conditions similar to the ones for data in Fig. 5A, but without an ATP regeneration system, myosin-X exhausts the 2 µM ATP of the buffer, leading to motor stalling and the stop of the barbed end elongation. (right) schematics presenting a myosin-X reaching the barbed end of an actin filament. When one motor domain is within ∼36 nm from the barbed end, the other motor domain may interfere with the arrival of an actin monomer or the departure of the last actin subunit while attempting to find its next actin binding site.

**Supp. Fig. S15.**
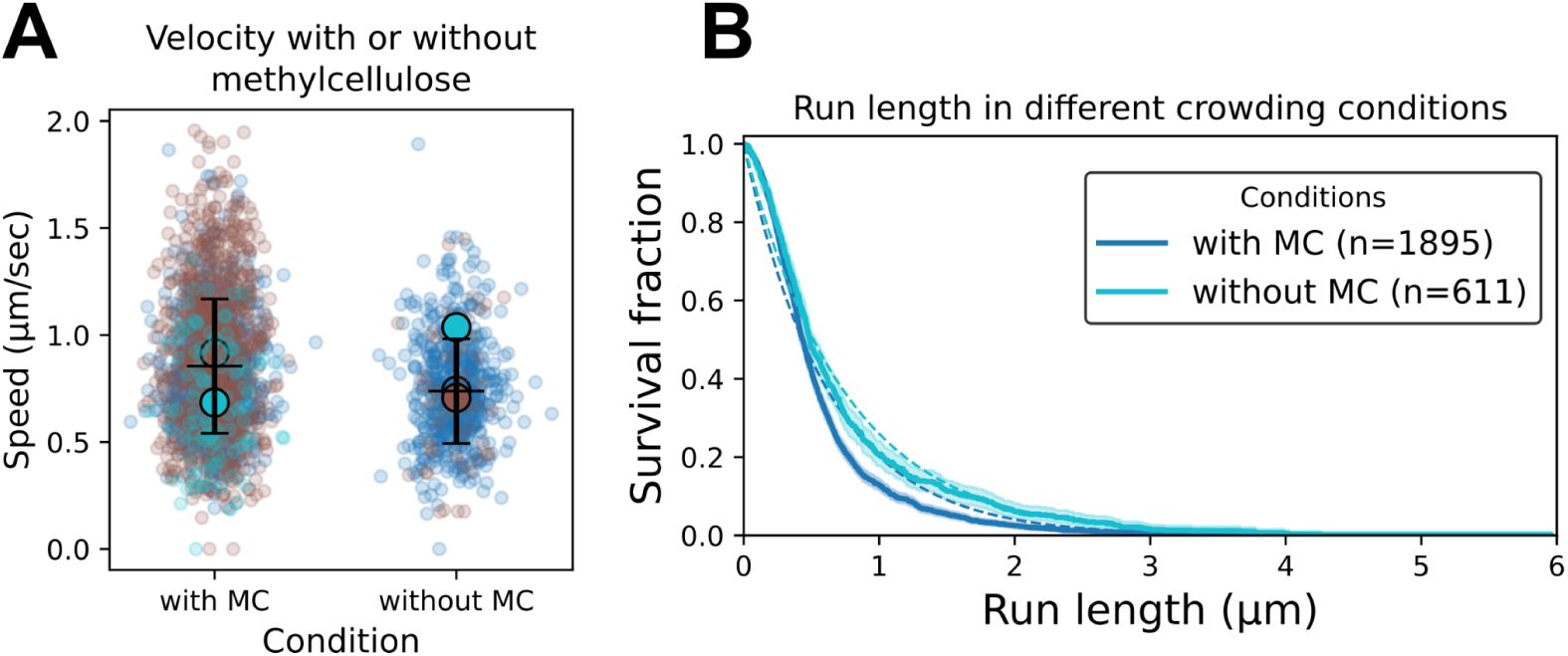
Effect of methylcellulose on the walking behaviour of myosin-X, at 200 µM ATP. (A) Velocity with (n = 1907, 0.85 ± 0.31 µm/s ) and without methylcellulose (n = 619, 0.74 ± 0.24 µm/s). (B) Kaplan-Meier survival curves of run length with (0.62 µm, 95% CI: 0.59-0.66 µm) and without methylcellulose (0.74 µm, 95% CI: 0.67-0.82 µm). Data from three independent experiments.

## Data Availability

The data are available from the corresponding author upon reasonable request.

## Acknowledgements

We thank the Romet/Jegou lab members for stimulating discussions, in particular Cécile Leduc and Hugo Wioland for their insightful comments on the manuscript.

## Author contributions

Wouter Kools, Trayana Hristova, Arliette Carlier and Sara Faour performed and analysed experiments. Elena-Maria Sirkia purified myosin-X. Ludivine Chaix and Antony Lee supplied reagents and optimized protocols. Anne Houdusse provided key protein resources. Anne Houdusse, Guillaume Romet-Lemonne, Pascal Martin and Antoine Jégou acquired funding. Guillaume Romet-Lemonne and Antoine Jégou designed and supervised the project. Antoine Jégou, and Wouter Kools wrote the manuscript. All authors reviewed the manuscript.

## Materials and Methods

### Proteins

Alpha skeletal muscle actin (UniProt P68135) was purified from rabbit muscle acetone powder following the initial protocol from Spudich and Watt (Spudich and Watt, 1971). Briefly, rabbit muscle acetonic powder was obtained from fresh rabbit muscles which were chopped then washed by 2 rounds of resuspension in Extraction Buffer (500 mM KCl, 50 mM KHCO_3_) and centrifugation (4000 g, 10 minutes, 4°C), then two rounds of resuspension in water whose pH was adjusted with NaCO_3_ to 8.6 and centrifugation (4000 g, 10 minutes, 4°C). The final pellets were resuspended by manual agitation, frozen and blended in −20°C acetone. The blended frozen muscles were filtered and the operation was repeated twice. The resulting muscle paste was spread out and dried overnight at room temperature. The powder was recovered and stored at −20°C until use. Acetonic powder was resuspended in X Buffer (2 mM Tris pH 7.8, 0.5 mM ATP, 0.1 mM CaCl_2_, 1 mM DTT, 0.01 % NaN_3_) and centrifuged (40,000 g, 45 minutes, 4°C). The supernatant was collected and filtered through glasswool. KCl was added to a final concentration of 3.3 M to remove contaminants and the solution centrifuged and filtered as before. The supernatant was then dialyzed overnight against 32 supernatant volumes of Dialysis Buffer (2 mM Tris pH 7.8, 1 mM MgCl_2_, 1 mM DTT), which brought the KCl concentration to 0.1 M. KCl was then added to a final concentration of 0.8 M, and the solution was incubated under agitation for 1h30 at 4°C, then ultracentrifuged at 100,000 g, 3h30, 4°C. The resulting pellet was resuspended with a potter in X buffer supplemented with 40 mM KCl and 2 mM MgCl_2_ and incubated overnight at 4°C. KCl was then added to increase its concentration to 0.8 M. The solution was further incubated under agitation 1h30 at 4°C then ultracentrifuged at 100,000 g, 3h30, 4°C. Repeating those steps as described here allows for a better detachment of undesired actin binding proteins. The resulting pellet was resuspended using a potter into G-buffer (2 mM Tris pH 7.8, 0.1 mM CaCl_2_, 0.01 % NaN_3_, 0.2 mM ATP, 1 mM DTT), and dialysed against the same buffer to depolymerize filaments at 4°C for 3 days. The solution was recovered, ultracentrifuged (400,000 g, 45 minutes, 4◦ C) and injected into an equilibrated Superdex 200 Hiload column (Cytiva) and actin was eluted with G-buffer into 1 mL fractions. Actin-containing fractions were identified by absorption at 290 nm, validated by electrophoresis and pooled. The left-most fractions were excluded as they might contain actin oligomers. The concentration was determined by measuring the optical density at 290 nm. Actin was stored on ice for up to 8 weeks, or flash frozen into liquid nitrogen and stored at −70◦ C.

Actin was fluorescently labeled on the surface accessible lysines using Alexa Fluor 488, 568, or 647 NHS ester (Thermo Fisher Scientific) as follows. Actin was dialysed overnight in a modified F-buffer (20 mM PIPES pH 6.9, 100 mM KCl, 0.2 mM ATP, 0.2 mM CaCl_2_) for polymerization. The resulting filaments were incubated with a 5x excess of fluorophore for 2h at room temperature on a rotating wheel, then ultracentrifuged at 350,000 g, 30 minutes, at room temperature. The resulting pellet was resuspended in G-buffer with a potter and the actin was left to depolymerize on ice for 2 hours. A new round of polymerization was induced by adding 400 mM KCl and 2 mM MgCl_2_ for 1 hour at room temperature. The filaments were ultracentrifuged 350,000 g, 30 minutes, at room temperature, and resuspended in G-buffer with a potter. The solution was dialysed overnight against G-buffer, then ultracentrifuged 350,000 g, 30 minutes, at room temperature. The concentration and labelling fraction were determined by measuring the optical density at 280 and the wavelength of the Alexa dyes. Final labelling fraction was between 15 to 45%, depending on the fluorophore, and used at 10% unless otherwise stated in the main text.

Spectrin-actin seeds were purified from human erythrocytes as follows. Human erythrocytes were first washed by repeated cycles of centrifugation (3,000 g, 15 minutes, at 4°C) and pellet resuspension in PBS buffer with EDTA (5 mM NaPO_4_ pH 7.7, 150 mM NaCl, 1 mM EDTA). Erythrocytes were then lysed in a low ionic strength buffer (5 mM NaPO_4_ pH 7.7, 1 mM PMSF) to turn them into cell ghosts. Cell ghosts were then washed and concentrated by 3 cycles of centrifugation (45,000 g, 15 minutes, at 4°C) and pellet resuspension in Washing Buffer (5 mM NaPO_4_ pH 7.7, 0.1 mM PMSF). The final pellet was resuspended and incubated at 37°C for 40 min with occasional mixing. Cells were then ultracentrifuged at 400,000 g, for 1 hour, at 4°C, to remove membrane residues. The supernatant was complemented with 2 mM DTT and protease inhibitors, and an equal volume of ice-cold glycerol was added. The concentration of functional spectrin-actin seeds was determined using a pyrene actin assay, to measure the concentration of growing filament barbed ends, using a spectro-fluorometer (Xenus instrument from SAFAS (Monaco); 10% labeled pyrene actin, excitation 366 nm, emission 407 nm; at room temperature). Spectrin-actin seed solution was then aliquoted and stored at −20°C until used (stable for months).

Recombinant human profilin1 (Uniprot: P07737) was expressed in E. coli (BL21 DE3 Star, Thermo Fischer) at 37°C for 2.5 hours. After centrifugation at 2,000 g the pellet was resuspended in Lysis buffer (Tris 50 mM pH 7.3, 5 mM EGTA, 0.1 mM EDTA, 50 mM KCl, 10 mM DTT, 8 M urea, 0.1% Tween−20, 1 mM PMSF, inhibitors). After sonication, the lysate was centrifuged at 186,000 g, 30 minutes, 4°C. The supernatant was dialysed overnight at 4°C against the Dialysis Buffer (Tris 50 mM pH 7.3, 1 mM EGTA, 0.1 mM EDTA, 50 mM KCl, 1 mM DTT). The protein solution was loaded onto a poly-L-proline column, washed with 3 column volumes of Washing Buffer (mixture of 3 volumes of Dialysis Buffer + 1 volume Elution Buffer), and eluted with Elution Buffer (Tris 50 mM pH 7.3, 5 mM EGTA, 0.1 mM EDTA, 50 mM KCl, 10 mM DTT, 8 M urea). Peak fractions were pooled, dialysed overnight at 4C against Dialysis buffer, concentrated with Vivaspin (cut-off 10kDa), and dialysed overnight at 4°C against Conservation Buffer (Tris 10 mM pH 7.5, 50 mM KCl, 1 mM DTT). The concentration was determined by absorption at 280 nm using the calculated extinction coefficient (17,020 M^-1^.cm^-1^). The protein solution was aliquoted, flash-frozen and stored at −70°C.

Two different myosin-X constructs were produced and purified: one fused to a mStayGold fluorescent protein, one without. Recombinant truncated human myosin-X (Uniprot Q9HD67, amino-acids 1 to 938) was followed by a 19 amino-acid linker (SEGGSGGSGGSGGSAASAA), a GCN4 leucine zipper to induce forced dimers (MKQLEDKVEELLSKNYHLENEVARLKKLVGE), a 5 amino-acid linker (GGSGGA) followed by a SnoopTag (KLGDIEFIKVN), a 4 amino-acid linker (KGGA), optionally an mStayGold fluorescent protein, a 6 amino-acid linker (SGTGGS) and a FLAG-tag, was expressed in the baculovirus/Sf9 insect cells expression system. Calmodulin from *Gallus gallus* was co-expressed on the same bacmid. The protein was purified using FLAG affinity chromatography followed by size exclusion chromatography, and stored at −20°C in 5 mM imidazole pH 7.0, 75 mM KCl, 10 mM DTT, 0.1 mM MgCl2, 0.5 mM EGTA, 50 μM MgADP and 50% glycerol. Mentioned myosin-X concentrations are for dimers.

### Buffers

All experiments were performed in standard F-buffer containing 5 mM Tris-HCl at pH 7.4, 1 mM MgCl_2_, 0.2 mM EGTA, 10 mM DTT, 1 mM DABCO, 50 mM KCl, supplemented with various concentrations of Mg-ATP and/or Mg-ADP. In the absence of ADP, we used an ATP regeneration system (10 mM creatine phosphate and 23 µg/mL (∼8 U/mL) creatine kinase) to maintain a stable ATP concentration.

‘Open’ chambers experiments were performed using F-buffer supplemented with 0.2 % methylcellulose (4,000 cP at 2%, 25°C, M0512, SIgma).

### Data acquisition

Experiments were performed using a Nikon TiE inverted microscope equipped with a TIRF CFI Apo 100× 1.49 NA oil-immersion objective, a Kinetix22 sCMOS camera (Photometrics), and a Total Internal Reflection Fluorescence (TIRF) illumination setup (iLAS2, Gataca Systems) with 100 mW 488-, 561-, and 642-nm tunable lasers. The temperature was maintained in chamber assay at 25 (± 0.2)°C using a collar objective heater (Okolab). The setup was controlled using micromanager (Edelstein et al., 2014).

### Microfluidics experiments

Microfluidics experiments were conducted in polydimethylsiloxane (PDMS; Sylgard) chambers based on the original protocol from (Jégou et al., 2011), described in detail in (Wioland et al., 2022). First, spectrin-actin seeds were attached to the glass surface by flowing in a solution containing 12 pM in F-buffer, for 1 minute. Next, the surface was passivated by exposing it to a solution containing 1 mg/mL PLL-PEG, for at least 30 minutes at room temperature in a humidity chamber. Typically, anchored actin filaments were polymerized from spectrin-actin seeds by flowing in 1 μM 10% Alexa Fluor 568–labeled G-actin for 5 to 10 min.

### ‘Open’ chamber experiments

We prepared ‘open’ chambers by melting parafilm stripes sandwiched in between cleaned coverslips. This creates chambers of around 10 µL. Chambers are either passivated with PLL-PEG (Fig. 4), as described above, or with 1 mg/mL PEG-Silane in 95% ethanol pH 2 dried on the coverslip at 70°C and thoroughly rinsed with pure water (Fig. 1). As a final step, the chamber is then extensively rinsed with an F-buffer.

### Attached filament experiments

To study the effect of methylcellulose on myosin-X (Supp. Fig. S15), we performed experiments with anchored actin filaments. The surface of an open chamber, passivated with 1:1000 biotin-PEG-silane:PEG-silane, was further functionalized with 10 µg/mL neutrAvidin for 1 min and washed with F-buffer. 0.5 µM pre-polymerised actin filaments (10% actin-Alexa488, 10% biotinylated actin) were injected and left to bind for 5 minutes. After rinsing, 15 nM myosin-X was injected into the chamber. Images were acquired in stream acquisition in TIRF mode with an exposure time of 100 ms. Before the myosin stream acquisition, a shorter stream acquisition of the actin channel was taken to exclude poorly attached filaments. Myosin-X movement was analysed using the FIJI plugin TrackMate (Ershov et al., 2022).

### FRAP data acquisition

A pre-defined ROI of 60x60 pixels was positioned over myosin-X induced bundles, such that either the tip or the full bundle was photobleached. Laser power was 100%, and the photobleaching duration was 2s. Images were then taken every 100 ms for the first 12 s, then every 5 s.

### Data analysis

#### Quantification of bundle size

The brightest 1 µm-long region of a bundle was normalised to the mean intensity of single filaments acquired on the same day in non-bundling conditions.

#### Quantification of the number of myosin-X at bundle tips

The maximum intensity in a 12x12 pixels box was normalised to that of single myosins acquired on the same day.

### FRAP analysis

The mean intensity in a box of 10x10 pixels around the tip was measured over time and normalised to the mean intensity before photobleaching. When the entire bundle was photobleached, a segmented line was drawn along the bundle and the intensity profile was extracted over time, normalised to the mean intensity before photobleaching.

### Myosin-X velocity

Kymographs from acquired movies were created by drawing lines along actin filaments or bundles using FiJi/ImageJ. Velocity was computed by measuring the distance traveled by each individual myosin divided by the duration of the run.

To obtain the ATP-dependent and ATP-independent rates of the ATPas cycle provided in the main text, myosin-X velocity as a function of ATP concentration (Supp. Fig. S6A) was fitted by the following formula: V = d.k_off,ADP_.k_on,ATP_.[ATP]/(k_off,ADP_+k_onATP_.[ATP]), assuming a step size d of 36 nm for myosin-X on single actin filaments, according to previously published results (Ropars et al., 2016; Sun et al., 2010). Errors are standard deviations. The fit shown in Supp Fig. S6A is a simple Michaelis-Menten saturation curve.

### Processivity and run length of myosin-X

To derive myosin-X processivity and run length, the fraction of processive myosins as a function of time or traveled distance is computed for detected myosins that can be tracked at least 3 consecutive frames. Survival fractions were constructed using the Kaplan-Meier method, using the ‘lifelines’ package in python. Censoring was applied when a myosin run was interrupted upon reaching a filament barbed end. The error bars represent the standard error calculated by the method of Greenwood.

All fits were done using the least square minimization procedure of the ‘curve_fit’ function from the Scipy python package. Reported errors are standard deviations, unless otherwise stated.

## References

Aramaki S, Mayanagi K, Jin M, Aoyama K, Yasunaga T. 2016. Filopodia Formation by Cross-linking of F-actin with Fascin in Two Different Binding Manners. Cytoskeleton . DOI: 10.1002/cm.21309, PMID: 27169557

Arsenault ME, Sun Y, Bau HH, Goldman YE. 2009. Using electrical and optical tweezers to facilitate studies of molecular motors. Physical Chemistry Chemical Physics 11:4834–4839. DOI: 10.1039/b821861g, PMID: 19506758

Baboolal TG, Mashanov GI, Nenasheva TA, Peckham M, Molloy JE. 2016. A Combination of Diffusion and Active Translocation Localizes Myosin 10 to the Filopodial Tip*. The Journal of biological chemistry 291:22373–22385. DOI: 10.1074/jbc.M116.730689

Bagès C, Chabanon M, Kools W, Dos Santos T, Pagès R, Sirkia ME, Leduc C, Houdusse A, Jégou A, Romet-Lemonne G, Wioland H. 2025. Probing protein-protein interactions with drag flow: a case study of F-actin and tropomyosin. The European physical journal. E, Soft matter 48:49. DOI: 10.1140/epje/s10189-025-00509-z, PMID: 40802217

Bao J, Huck D, Gunther LK, Sellers JR, Sakamoto T. 2013. Actin structure-dependent stepping of myosin 5a and 10 during processive movement. PloS one 8:e74936. DOI: 10.1371/journal.pone.0074936, PMID: 24069366

Berg JS, Cheney RE. 2002. Myosin-X is an unconventional myosin that undergoes intrafilopodial motility. Nature cell biology 4:246–250. DOI: 10.1038/ncb762, PMID: 11854753

Berg JS, Derfler BH, Pennisi CM, Corey DP, Cheney RE. 2000. Myosin-X, a novel myosin with pleckstrin homology domains, associates with regions of dynamic actin. Journal of Cell Science 113 Pt 19:3439–3451. DOI: 10.1242/jcs.113.19.3439, PMID: 10984435

Bieling P, Telley IA, Surrey T. 2010. A minimal midzone protein module controls formation and length of antiparallel microtubule overlaps. Cell 142:420–432. DOI: 10.1016/j.cell.2010.06.033, PMID: 20691901

Blake TCA, Gallop JL. 2023. Filopodia In Vitro and In Vivo. Annual review of cell and developmental biology. DOI: 10.1146/annurev-cellbio-020223-025210, PMID: 37406300

Bohil AB, Robertson BW, Cheney RE. 2006. Myosin-X is a molecular motor that functions in filopodia formation. Proceedings of the National Academy of Sciences of the United States of America 103:12411–12416. DOI: 10.1073/pnas.0602443103, PMID: 16894163

Bornschlögl T. 2013. How filopodia pull: what we know about the mechanics and dynamics of filopodia. Cytoskeleton 70:590–603. DOI: 10.1002/cm.21130, PMID: 23959922

Caporizzo MA, Fishman CE, Sato O, Jamiolkowski RM, Ikebe M, Goldman YE. 2018. The Antiparallel Dimerization of Myosin X Imparts Bundle Selectivity for Processive Motility. Biophysical journal 114:1400–1410. DOI: 10.1016/j.bpj.2018.01.038, PMID: 29590597

Cheffings TH, Burroughs NJ, Balasubramanian MK. 2016. Actomyosin Ring Formation and Tension Generation in Eukaryotic Cytokinesis. Current biology: CB 26:R719–R737. DOI: 10.1016/j.cub.2016.06.071, PMID: 27505246

Claessens MMAE, Bathe M, Frey E, Bausch AR. 2006. Actin-binding proteins sensitively mediate F-actin bundle stiffness. Nature materials 5:748–753. DOI: 10.1038/nmat1718

Edelstein AD, Tsuchida MA, Amodaj N, Pinkard H, Vale RD, Stuurman N. 2014. Advanced methods of microscope control using μManager software. Journal of biological methods 1. DOI: 10.14440/jbm.2014.36, PMID: 25606571

Ershov D, Phan M-S, Pylvänäinen JW, Rigaud SU, Le Blanc L, Charles-Orszag A, Conway JRW, Laine RF, Roy NH, Bonazzi D, Duménil G, Jacquemet G, Tinevez J-Y. 2022. TrackMate 7: integrating state-of-the-art segmentation algorithms into tracking pipelines. Nature Methods 19:829–832. DOI: 10.1038/s41592-022-01507-1, PMID: 35654950

Faix J, Rottner K. 2006. The making of filopodia. Current opinion in cell biology 18:18–25. DOI: 10.1016/j.ceb.2005.11.002, PMID: 16337369

Fitz GN, Weck ML, Bodnya C, Perkins OL, Tyska MJ. 2023. Protrusion growth driven by myosin-generated force. Developmental Cell 58:18–33.e6. DOI: 10.1016/j.devcel.2022.12.001, PMID: 36626869

Gong R, Jiang F, Moreland ZG, Reynolds MJ, los Reyes SE de, Gurel P, Shams A, Heidings JB, Bowl MR, Bird JE, Alushin GM. 2022. Structural basis for tunable control of actin dynamics by myosin-15 in mechanosensory stereocilia. Science Advances 8:eabl4733. DOI: 10.1126/sciadv.abl4733

Gong R, Reynolds MJ, Carney KR, Hamilton K, Bidone TC, Alushin GM. 2025. Fascin structural plasticity mediates flexible actin bundle construction. Nature structural & molecular biology 32:940–952. DOI: 10.1038/s41594-024-01477-2, PMID: 39833469

Holland SM, Gallo G. 2023. Actin cytoskeletal dynamics do not impose an energy drain on growth cone bioenergetics. Journal of cell science. DOI: 10.1242/jcs.261356, PMID: 37534394

Homma K, Ikebe M. 2005. Myosin X Is a High Duty Ratio Motor*. The Journal of biological chemistry 280:29381–29391. DOI: 10.1074/jbc.M504779200

Houdusse A, Titus MA. 2021. The many roles of myosins in filopodia, microvilli and stereocilia. Current biology: CB 31:R586–R602. DOI: 10.1016/j.cub.2021.04.005, PMID: 34033792

Jacquemet G, Hamidi H, Ivaska J. 2015. Filopodia in cell adhesion, 3D migration and cancer cell invasion. Current opinion in cell biology 36:23–31. DOI: 10.1016/j.ceb.2015.06.007, PMID: 26186729

Jacquemet G, Stubb A, Saup R, Miihkinen M, Kremneva E, Hamidi H, Ivaska J.\ 2019. Filopodome Mapping Identifies p130Cas as a Mechanosensitive Regulator of Filopodia Stability. Current biology: CB 29:202–216.e7. DOI: 10.1016/j.cub.2018.11.053, PMID: 30639111

Jégou A, Niedermayer T, Orbán J, Didry D, Lipowsky R, Carlier M-F, Romet-Lemonne G. 2011. Individual actin filaments in a microfluidic flow reveal the mechanism of ATP hydrolysis and give insight into the properties of profilin. PLoS biology 9:e1001161. DOI: 10.1371/journal.pbio.1001161, PMID: 21980262

Knight PJ, Thirumurugan K, Xu Y, Wang F, Kalverda AP, Stafford WF 3rd, Sellers JR, Peckham M. 2005. The predicted coiled-coil domain of myosin 10 forms a novel elongated domain that lengthens the head. The Journal of Biological Chemistry 280:34702–34708. DOI: 10.1074/jbc.M504887200, PMID:16030012

Kučera O, Siahaan V, Janda D, Dijkstra SH, Pilátová E, Zatecka E, Diez S, Braun M, Lansky Z. 2021. Anillin propels myosin-independent constriction of actin rings. Nature communications 12:4595. DOI: 10.1038/s41467-021-24474-1, PMID: 34321459

Lansky Z, Braun M, Lüdecke A, Schlierf M, ten Wolde PR, Janson ME, Diez S. 2015. Diffusible crosslinkers generate directed forces in microtubule networks. Cell 160:1159–1168. DOI: 10.1016/j.cell.2015.01.051, PMID: 25748652

Leduc C, Padberg-Gehle K, Varga V, Helbing D, Diez S, Howard J. 2012. Molecular crowding creates traffic jams of kinesin motors on microtubules. Proceedings of the National Academy of Sciences of the United States of America 109:6100–6105. DOI: 10.1073/pnas.1107281109, PMID: 22431622

Lehtimäki JI, Rajakylä EK, Tojkander S, Lappalainen P. 2021. Generation of stress fibers through myosin-driven reorganization of the actin cortex. eLife 10. DOI: 10.7554/eLife.60710, PMID: 33506761

Leijnse N, Barooji YF, Arastoo MR, Sønder SL, Verhagen B, Wullkopf L, Erler JT, Semsey S, Nylandsted J, Oddershede LB, Doostmohammadi A, Bendix PM. 2022. Filopodia rotate and coil by actively generating twist in their actin shaft. Nature communications 13:1636. DOI: 10.1038/s41467-022-28961-x, PMID: 35347113

Leijnse N, Oddershede LB, Bendix PM. 2015. Helical buckling of actin inside filopodia generates traction. Proceedings of the National Academy of Sciences of the United States of America 112:136–141. DOI: 10.1073/pnas.1411761112, PMID: 25535347

Lenz M, Gardel ML, Dinner AR. 2012. Requirements for contractility in disordered cytoskeletal bundles. New journal of physics 14:033037. DOI: 10.1088/1367-2630/14/3/033037, PMID: 23155355

Lu Q, Ye F, Wei Z, Wen Z, Zhang M. 2012. Antiparallel coiled-coil-mediated dimerization of myosin X. Proceedings of the National Academy of Sciences of the United States of America 109:17388–17393. DOI: 10.1073/pnas.1208642109, PMID: 23012428

Miihkinen M, Grönloh MLB, Popović A, Vihinen H, Jokitalo E, Goult BT, Ivaska J, Jacquemet G. 2021. Myosin-X and talin modulate integrin activity at filopodia tips. Cell reports 36:109716. DOI: 10.1016/j.celrep.2021.109716, PMID: 34525374

Moreland ZG, Jiang F, Aguilar C, Barzik M, Gong R, Behnammanesh G, Park J, Shams A, Faaborg-Andersen C, Werth JC, Harley R, Sutton DC, Heidings JB, Cole SM, Parker A, Morse S, Wilson E, Takagi Y, Sellers JR, Brown SDM, Friedman TB, Alushin GM, Bowl MR, Bird JE. 2025. Myosin-based nucleation of actin filaments contributes to stereocilia development critical for hearing. Nature Communications 16:947. DOI: 10.1038/s41467-025-55898-8, PMID: 39843411

Murrell M, Oakes PW, Lenz M, Gardel ML. 2015. Forcing cells into shape: the mechanics of actomyosin contractility. Nature reviews. Molecular cell biology. DOI: 10.1038/nrm4012, PMID: 26130009

Murrell MP, Gardel ML. 2012. F-actin buckling coordinates contractility and severing in a biomimetic actomyosin cortex. Proceedings of the National Academy of Sciences 109:20820–20825. DOI: 10.1073/pnas.1214753109

Nagy S, Ricca BL, Norstrom MF, Courson DS, Brawley CM, Smithback PA, Rock RS. 2008. A myosin motor that selects bundled actin for motility. Proceedings of the National Academy of Sciences of the United States of America 105:9616–9620. DOI: 10.1073/pnas.0802592105, PMID: 18599451

Nguyen QQ, Zhou Y, Cheng MS, Qin X, Cheng HCM, Liu X, Sweeney HL, Park H. 2023. The Antiparallel Coiled-Coil Domain Allows Multiple Forward Step Sizes of Myosin X. Journal of Physical Chemistry Letters 4914–4922. DOI: 10.1021/acs.jpclett.3c00512, PMID: 37202741

Pernier J, Kusters R, Bousquet H, Lagny T, Morchain A, Joanny J-F, Bassereau P, Coudrier E. 2019. Myosin 1b is an actin depolymerase. Nature communications 10:5200. DOI: 10.1038/s41467-019-13160-y, PMID: 31729365

Pokrant T, Hein JI, Körber S, Disanza A, Pich A, Scita G, Rottner K, Faix J. 2023. Ena/VASP clustering at microspike tips involves lamellipodin but not I-BAR proteins, and absolutely requires unconventional myosin-X. Proceedings of the National Academy of Sciences of the United States of America 120:e2217437120. DOI: 10.1073/pnas.2217437120, PMID: 36598940

Popović A, Miihkinen M, Ghimire S, Saup R, Grönloh MLB, Ball NJ, Goult BT, Ivaska J, Jacquemet G. 2023. Myosin-X recruits lamellipodin to filopodia tips. Journal of cell science 136. DOI: 10.1242/jcs.260574, PMID: 36861887

Ropars V, Yang Z, Isabet T, Blanc F, Zhou K, Lin T, Liu X, Hissier P, Samazan F, Amigues B, Yang ED, Park H, Pylypenko O, Cecchini M, Sindelar CV, Sweeney HL, Houdusse A. 2016. The myosin X motor is optimized for movement on actin bundles. Nature communications 7:12456. DOI: 10.1038/ncomms12456, PMID: 27580874

Sakamoto T, Webb MR, Forgacs E, White HD, Sellers JR. 2008. Direct observation of the mechanochemical coupling in myosin Va during processive movement. Nature 455:128–132. DOI: 10.1038/nature07188, PMID: 18668042

Sato O, Jung HS, Komatsu S, Tsukasaki Y, Watanabe TM, Homma K, Ikebe M. 2017. Activated full-length myosin-X moves processively on filopodia with large steps toward diverse two-dimensional directions. Scientific reports 7:44237. DOI: 10.1038/srep44237, PMID: 28287133

Schuler M-H, Lewandowska A, Caprio GD, Skillern W, Upadhyayula S, Kirchhausen T, Shaw JM, Cunniff B. 2017. Miro1-mediated mitochondrial positioning shapes intracellular energy gradients required for cell migration. Molecular biology of the cell 28:2159–2169. DOI: 10.1091/mbc.E16-10-0741, PMID: 28615318

Sebé-Pedrós A, Grau-Bové X, Richards TA, Ruiz-Trillo I. 2014. Evolution and classification of myosins, a paneukaryotic whole-genome approach. Genome Biology and Evolution 6:290–305. DOI: 10.1093/gbe/evu013, PMID: 24443438

Spudich JA, Watt S. 1971. The regulation of rabbit skeletal muscle contraction. I. Biochemical studies of the interaction of the tropomyosin-troponin complex with actin and the proteolytic fragments of myosin. The Journal of biological chemistry 246:4866–4871. DOI: 10.1016/S0021-9258(18)62016-2, PMID: 4254541

Sun Y, Sato O, Ruhnow F, Arsenault ME, Ikebe M, Goldman YE. 2010. Single-molecule stepping and structural dynamics of myosin X. Nature structural & molecular biology 17:485–491. DOI: 10.1038/nsmb.1785, PMID: 20364131

Svitkina TM, Bulanova EA, Chaga OY, Vignjevic DM, Kojima S-I, Vasiliev JM, Borisy GG. 2003. Mechanism of filopodia initiation by reorganization of a dendritic network. The Journal of cell biology 160:409–421. DOI: 10.1083/jcb.200210174, PMID: 12566431

Takagi Y, Farrow RE, Billington N, Nagy A, Batters C, Yang Y, Sellers JR, Molloy JE. 2014. Myosin-10 produces its power-stroke in two phases and moves processively along a single actin filament under low load. Proceedings of the National Academy of Sciences of the United States of America 111:E1833–42. DOI: 10.1073/pnas.1320122111, PMID: 24753602

Tantama M, Martínez-François JR, Mongeon R, Yellen G. 2013. Imaging energy status in live cells with a fluorescent biosensor of the intracellular ATP-to-ADP ratio. Nature Communications 4:2550. DOI: 10.1038/ncomms3550, PMID: 24096541

Thoresen T, Lenz M, Gardel ML. 2011. Reconstitution of Contractile Actomyosin Bundles. Biophysical journal 100:2698–2705. DOI: 10.1016/j.bpj.2011.04.031

Tokuo H, Ikebe M. 2004. Myosin X transports Mena/VASP to the tip of filopodia. Biochemical and biophysical research communications 319:214–220. DOI: 10.1016/j.bbrc.2004.04.167, PMID: 15158464

Tokuo H, Mabuchi K, Ikebe M. 2007. The motor activity of myosin-X promotes actin fiber convergence at the cell periphery to initiate filopodia formation. The Journal of cell biology 179:229–238. DOI: 10.1083/jcb.200703178, PMID: 17954606

Truong Quang BA, Peters R, Cassani DAD, Chugh P, Clark AG, Agnew M, Charras G, Paluch EK. 2021. Extent of myosin penetration within the actin cortex regulates cell surface mechanics. Nature communications 12:6511. DOI: 10.1038/s41467-021-26611-2, PMID: 34764258

Varga V, Leduc C, Bormuth V, Diez S, Howard J. 2009. Kinesin-8 motors act cooperatively to mediate length-dependent microtubule depolymerization. Cell 138:1174–1183. DOI: 10.1016/j.cell.2009.07.032, PMID: 19766569

Vignaud T, Copos C, Leterrier C, Toro-Nahuelpan M, Tseng Q, Mahamid J, Blanchoin L, Mogilner A, Théry M, Kurzawa L. 2021. Stress fibres are embedded in a contractile cortical network. Nature materials 20:410–420. DOI: 10.1038/s41563-020-00825-z, PMID: 33077951

Wioland H, Ghasemi F, Chikireddy J, Romet-Lemonne G, Jégou A. 2022. Using microfluidics and fluorescence microscopy to study the assembly dynamics of single actin filaments and bundles. Journal of visualized experiments: JoVE. DOI: 10.3791/63891

Wioland H, Suzuki E, Cao L, Romet-Lemonne G, Jegou A. 2020. The advantages of microfluidics to study actin biochemistry and biomechanics. Journal of muscle research and cell motility 41:175–188. DOI: 10.1007/s10974-019-09564-4, PMID: 31749040

Wollrab V, Belmonte JM, Baldauf L, Leptin M, Nédeléc F, Koenderink GH. 2018. Polarity sorting drives remodeling of actin-myosin networks. Journal of cell science 132. DOI: 10.1242/jcs.219717, PMID: 30404824

Zemel A, Mogilner A. 2009. Motor-induced sliding of microtubule and actin bundles. Physical Chemistry Chemical Physics 11:4821–4833. DOI: 10.1039/b818482h, PMID: 19506757

